# Extensive multi-species hybridization between Leuciscidae minnow species

**DOI:** 10.1101/2025.07.14.664671

**Authors:** Amanda V. Meuser, Amy R. Pitura, S. Eryn McFarlane, Elizabeth G. Mandeville

**Author notes:** Corresponding author : Amanda V. Meuser Stockholm University Department of Zoology SE-106 91 Stockholm, Sweden.

## Abstract

Anthropogenic disturbances can disrupt ecosystems and alter species population dynamics. Interspecific hybridization is common between genetically related organisms, especially once reproductive barriers such as spatial isolation have been removed. We used genotyping-by-sequencing data to assess outcomes of hybridization between several Leuciscidae minnow species and to identify to what extent land use type and environmental variables influence the frequency of hybridization. We found that both two-species and multi-species hybridization was widespread; hybrids were sampled at all 25 sampling sites and made up almost 30% of all individuals sampled. While most species hybridized with at least one other sampled species, the amount of hybridization was variable. We used logistic regression to estimate the influence of anthropogenic disturbance on hybridization, and found significant but weak relationships between hybridization and environmental factors. This research improves our understanding of hybridization dynamics in species-rich clades like the Leuciscidae with low reproductive isolation, and points to the need for additional work to better understand predictors of hybridization in multi-species hybrid zones.

## 1 Introduction

Hybridization can be common between sympatric, closely related species, but hybrids are often difficult to detect. While some instances of hybridization can be identified morphologically, the extent of backcrossing and introgression is difficult to quantify and cryptic hybrids or backcrossed individuals can go entirely unnoticed (Arnold *et al*. 2003, Slager *et al*. 2020). Despite being difficult to detect, it is clear that hybridization has substantial but variable effects on biodiversity. While in some instances hybridization can lead to adaptive introgression or speciation, hybridization can also act as an avenue towards extinction and may produce a net loss of biodiversity through genetic or demographic swamping (Rhymer & Simberloff 1996, Todesco *et al*. 2016, Wolf *et al*. 2001). Hybridization and the proportion of hybrids in a population increase when disturbances alter prezygotic reproductive isolation (e.g. modifying breeding time, breeding location, or mate selection) or when disturbances create a scenario where hybrid individuals have higher fitness than parental species (Chunco 2014, Grabenstein & Taylor 2018, Todesco *et al*. 2016), making the interplay between disturbance and hybridization especially critical to understanding how hybridization affects biodiversity.

Anthropogenic disturbances to the natural environment alter species interactions and increase the number of hybrid zones formed (Hasselman *et al*. 2014, Jansson *et al*. 2007, Seehausen *et al*. 1997). Aquatic systems are particularly vulnerable to anthropogenic disturbances, such as eutrophication, pollution, sedimentation, deforestation, and channelization (Moss 2008, Moyle 1999, Rahel 2002, Scott & Helfman 2001). Despite both the clear link between hybridization and disturbance, as well as the prevalence of disturbance in aquatic systems, hybridization in fish remains incompletely described, even for very common species or clades. Leuciscid minnows are widespread in North America, abundant in small streams (Holm *et al*. 2022, Schultz 2003), and have been noted to naturally hybridize to some degree (Cooper 1980, Greenfield *et al*. 1973, Ross 1975, Ross & Cavender 1981). However, almost all previous work has identified hybridization between leuciscid species based on morphological characteristics alone (Holm *et al*. 2022). Intriguingly, recent research identified breeding behaviour as a mechanism regulating hybridization in minnows (Corush *et al*. 2020). The importance of breeding behaviour in promoting or inhibiting hybridization provides a clear potential mechanism for how anthropogenic disturbance might promote hybridization, specifically that nesting behaviours are likely affected by anthropogenic disturbance in this family of fishes. Increased hybridization has been noted in other species of fish which have had breeding behaviours affected by anthropogenic disturbances (Hasselman *et al*. 2014, Seehausen *et al*. 1997). However, while hybridization between leuciscid species has been described based on morphological identification, there has been little genomic work done until quite recently to confirm or quantify hybridization in leuciscids (but see Zbinden *et al*. 2023).

Morphological identifications of leuciscid minnows are difficult to make with confidence and require substantial expertise; intermediate hybrid phenotypes are even more difficult to assign accurately (Hubbs 1955). Hybrids are also not necessarily expected to have intermediate phenotypes. In many hybrid crosses across diverse clades, F_1_ hybrid phenotypes are quite variable and can be considerably biased towards one parent, making them — and especially backcrosses with either parental population — easy to miss entirely (Thompson *et al*. 2021). DNA sequence data can be used to estimate ancestry and detect introgression and backcrossing where morphological identification has failed or is insufficient (Arnold *et al*. 2003, Slager *et al*. 2020). Using genomic data is especially important when hybrid zones involve more than two species (Natola *et al*. 2022, Satokangas *et al*. 2023), as we suspected might be the case for leuciscid minnows.

The goal of this study was to investigate both the extent of hybridization between minnow species and how hybridization differed across areas of varied anthropogenic disturbance in Southern Ontario using genomic data. Specifically, we sampled and assessed hybridization with genomic data between these leuciscid fishes: creek chub (*Semotilus atromaculatus*), common shiner (*Luxilus cornutus*), central stoneroller (*Campostoma anomalum*), hornyhead chub (*Nocomis biguttatus*), western blacknose dace (*Rhinichthys obtusus*), striped shiner (*Luxilus chrysocephalus*), longnose dace (*Rhinichthys cataractae*), river chub (*Nocomis micropogon*), and rosyface shiner (*Notropis rubellus*). Then, we compared proportion of hybrids at each site with the land use and environmental data for the upstream watershed of each sampling site. We addressed two questions: (1) How do the frequency and type of hybrids vary between species pairs? and (2) To what extent do the type and degree of disturbance across the sampling locations influence hybridization outcomes?

## 2 Materials and Methods

### 2.1 Sampling and study sites

We performed the majority of sampling in summer 2022. We chose sampling sites that represented agriculturalized, urbanized, and low-disturbance areas. We first chose areas we knew to have minnows present, from either past sampling efforts or personal knowledge. Then, we used Google Earth (earth.google.com) to find additional sites in nearby agricultural or urbanized areas that were reasonably likely to contain our target species, based on the size and surrounding environment of each stream or lake. Finally, we consulted technical reports and online resources to confirm that our species of interest had previously been sampled from these sites. The sampled streams and rivers and the fauna that inhabit them have been greatly affected by the construction of roadways (Wallace *et al*. 2013), dredging and forced channelization Moss (2008), Ward-Campbell *et al*. (2017), and pollution from farming and other human activities (Liu *et al*. 2018, Moss 2008, Xu *et al*. 2020). Sites can be seen in Figure 1A.

**Figure 1:**
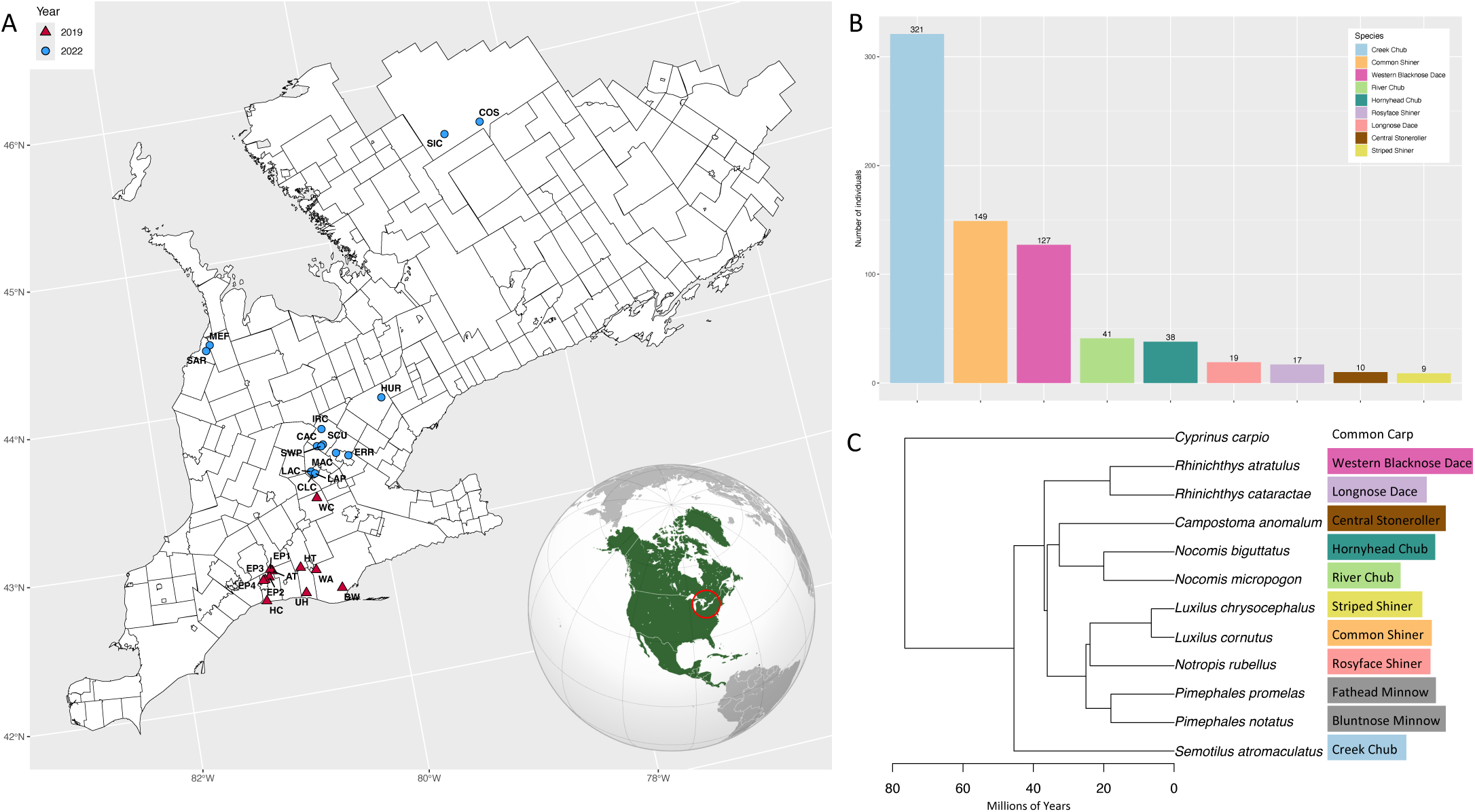
Sampling outcomes and phylogenetic relationship between study species, for individuals retained after data filtering. A. Map of sampling sites, coloured by sampling year, n = 25. B. Breakdown of sampled fish by species for both sampling years combined, identified by phenotype. n = 731. C. Phylogeny showing all 9 species of interest in this study, plus the two Pimephales species that were identified through DNA barcoding. This phylogeny was created with data from The Fish Tree of Life database, which was assembled using DNA sequence data from NCBI GenBank (Chang et al. 2019).

Additionally, we sampled fish from streams in Algonquin Provincial Park to represent lower-disturbance areas. While Algonquin is the oldest provincial park in Ontario, it still suffers impacts of anthropogenic disturbance in parts more frequented by tourists and has a long history of logging within park boundaries, starting in the early 1800s and contin- uing today (Helmers *et al*. 2016, McIntyre 2022). Chosen streams fall within the current Wilderness and Recreation/Utilization Zones, away from increased human activity within the Development and Natural Environment Zones; these sites include Simms Creek and Costello Creek. While we have aimed to choose sites surrounded by lower amounts of human disturbance, no area in Southern Ontario is truly undisturbed, and we deemed water bodies outside of Southern Ontario inappropriate as reference locations due to differences in geology and species distribution.

Some initial sampling was done by our collaborators in summer 2019 in connection with another study (Champagne *et al*. 2022, Gutgesell *et al*. 2025); creek chub, western blacknose dace, and common shiner samples were collected from agricultural streams across Southern Ontario. Additional creek chub samples were also collected from Costello Creek by our collaborators in summer 2022.

We sampled under the Ontario Ministry of Northern Development, Mines, Natural Resources and Forestry licence #1100698 and the University of Guelph’s Animal Utilization Protocol permit #4237. We caught minnows using both wire minnow traps and a straight seine net. We selected individual fish for sampling if they visually appeared to be members of one or more of the species of interest. We euthanized the sampled fish using an overdose of MS-222 (a 250 mg/L solution for 10 minutes), then cut either a pectoral fin from larger fish or the caudal fin from smaller fish and preserved both the fin and the body of the fish in two separate containers of 95% ethanol, for later DNA extraction. Samples collected by our collaborators in summers 2019 and 2022 were caught either using electrofishing or wire minnow traps, but were euthanized, dissected, and stored in the same fashion as our samples.

In total, we gathered 1213 fish across all field work efforts in 2019 and 2022 (Figure 1B).

We used the Ontario Watershed Information Tool (OWIT, ontario.ca/page/ontario-watershed-information-tool-owit) to gather exact water- shed characteristics and land cover information for each sampling site. We used the percent of total watershed area values for “Community Infrastructure”, “Agriculture and Undifferentiated Rural Land Use”, and “Sand Gravel Mine Tailings Extraction”as a proxy for representing how anthropogenically disturbed each site’s watershed is.

### 2.2 Genomic data collection

We extracted DNA from each of the samples using the QIAGEN DNeasy Blood & Tissue Kit (Qiagen, Inc.), according to the manufacturer’s instructions. We used a NanoDrop 8000 Spectrophotometer (ThermoFisher Scientific) to quantify DNA concentration and check that each sample was equal or greater to 20 ng/*µ*l. Following DNA extraction, we prepared genomic libraries using a reduced-representation genotyping by sequencing method (Parchman *et al*. 2012) to prepare samples for high-throughput sequencing. We digested DNA with EcoRI and MseI restriction enzymes, then ligated adaptors containing the adaptor sequence, barcode, cut site, and a protector base onto the fragments, before amplifying the fragments with PCR and annealing Illumina PCR primers onto either end (Parchman *et al*. 2012). We then size selected for fragments 200-300bp in length using the Sage Science PippinPrep. This fragment range captured the most frequent size of post-PCR DNA fragments produced by restriction enzyme digest, as confirmed by Tapestation results. The DNA samples were sequenced at The Centre for Applied Genomics in Toronto, Canada. There, single-end 100bp reads were sequenced with an Illumina NovaSeq 6000, using 2 lanes of an S1 flowcell.

### 2.3 Alignment and filtering of raw sequence data

We performed much of the computing necessary for this project via an allocation on the Digital Research Alliance of Canada’s high-performance computing cluster, Cedar, with a few analyses and data visualization in R using RStudio on a local computer (R Core Team 2021).

We demultiplexed raw FASTQ files using sabre (github.com/najoshi/sabre), with -m set to 2 to allow up to 2 barcode mismatches. Then we used the Burrows-Wheeler Alignment tool (BWA) (Li & Durbin 2009, version 0.7.17), with the BWA-MEM algorithm (Li 2013), to align the FASTQ files to the creek chub reference genome (Meuser *et al*. 2023). We used bcftools mpileup (Li & Durbin 2009, version 1.11) to identify variable loci; these were output into a BCF file which we converted to a VCF file using SAMtools (Li *et al*. 2009, version 1.16). We set the --max-missing parameter to 0.6 - i.e., at least 40% of individuals must have data at a locus to retain that locus and the --mac parameter to 3, to include only sites with Minor Allele Count greater than or equal to 3. Afterwards, we adjusted depth to keep only sites with depth values greater than or equal to the --minDP value of 3. Then, we generated an .imiss file reporting the missingness on a per-individual basis, created a file called lowDP.indiv with those that are missing greater than 95% of loci, and removed those individuals from the VCF file. Finally, we removed loci with a minor allele frequency of less than 1%, by setting the --maf parameter to 0.01. We also removed 69 individuals from library prep plate 13, as an issue with demultiplexing these individuals rendered them unusable. This left 731 individuals for further analysis in the main data set.

Additionally, we created three filtered VCF files for each pairwise comparison of creek chub (CC), common shiner (CS), and western blacknose dace (BND) – the three most abundant species in the data set – plus their interspecific hybrids. We did this to examined inter-source ancestry, as it can only be calculated between species pairs. We used VCFtools’ --keep function and input a list of all of the individuals to retain in the output file. These individuals were chosen based on genomic identifications of species from the output from ENTROPY that were classified as either parentals or hybrids of the two parental species.

### 2.4 Analysis of hybridization with **ENTROPY**

We used the software ENTROPY to assess admixture between each of the 9 sampled minnow species (Shastry *et al*. 2021). ENTROPY estimates admixture proportion, q (the predicted ancestry from either parental species), and inter-source ancestry, Q (the proportion of loci in each individual with an allele from each parental population) (Shastry *et al*. 2021). We used an R script (inputdataformat.R) (Shastry *et al*. 2021, R Core Team 2021) to create a multiple population genotype likelihood (MPGL), linear discriminant analysis k-means clustering (LDAK) files, and a point estimate file, from the VCF file. We first ran ENTROPY for all 800 individuals. As we had phenotypically identified 9 species, we ran 3 repetitions of K=1 through K=15, as K=9 may not necessarily be the best fitting model. We ran the program with 8GB of RAM and 150,000 steps total, including 100,000 burn-in steps, and thinned posterior chains to retain every 25th step following the burn-in. For the 3 data sets containing the pairwise species combinations, we ran 3 repetitions of ENTROPY for K=1 through K=3 as there should be 2 species in each data set. For these runs specifically, we also set the -Q parameter to 1, to estimate inter-source ancestry values for each individual. After running ENTROPY, we calculated the deviance information criterion (DIC) for each K value we submitted (Spiegelhalter *et al*. 2002), averaged across the 3 replicate chains, for each data set. We calculated this for the data by using first a custom shell script to loop over all results files and extract DIC from each HDF5 file using the estpost.entropy program. For each ENTROPY run, we also checked for proper mixing and convergence of the model by visually inspecting a subset of the MCMC chains.

Once we had the best value of K, we plotted the q values for all individuals, split by site (Fig. 2). We created triangle plots for K=2 models of pairwise combinations of creek chub, common shiner, and western blacknose dace, using both Q and q values (Fig.3A,C,E).

**Figure 2:**
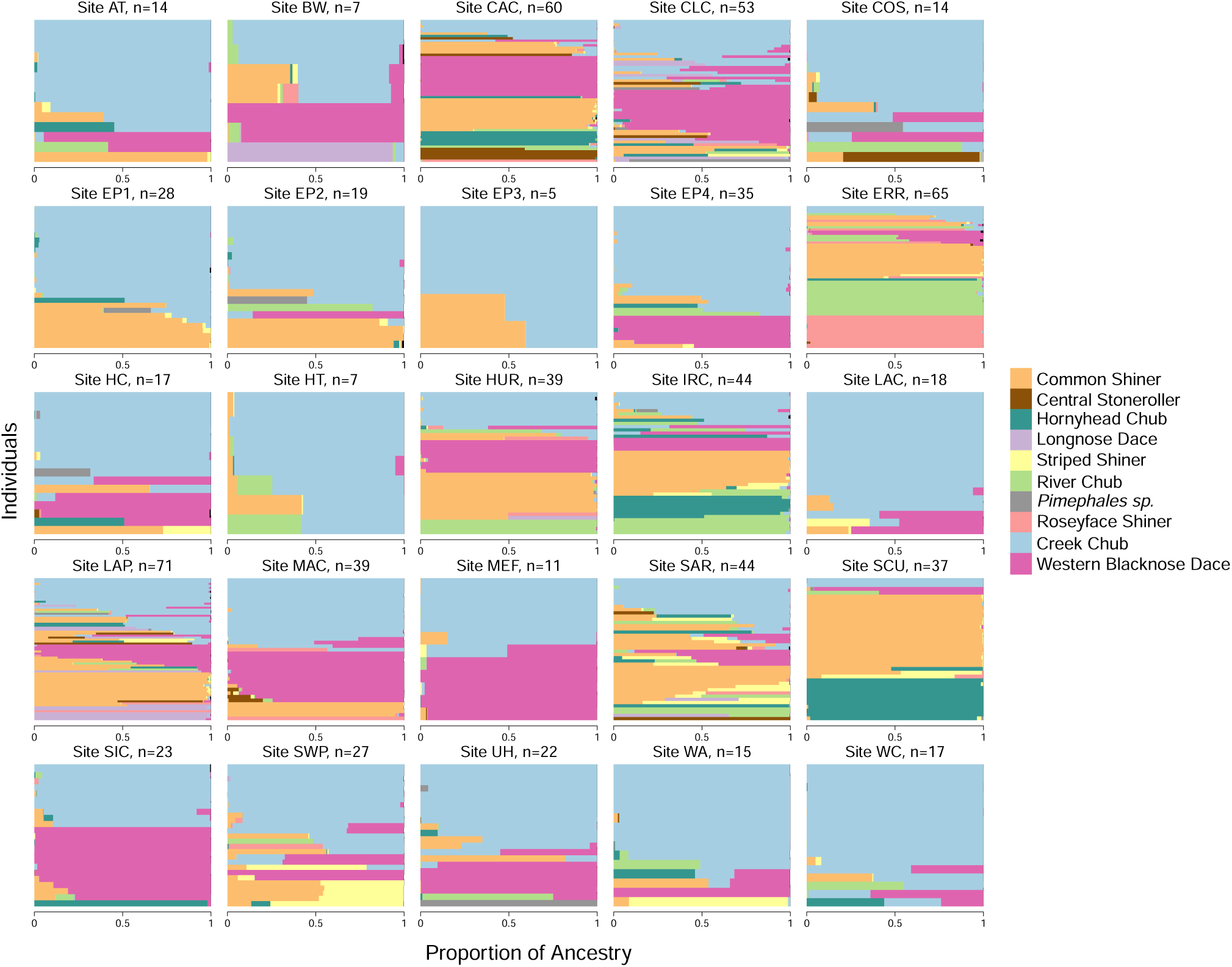
Results from ENTROPY are shown plotted by site. Individuals are represented by a horizontal bar, with proportion of ancestry from each population shown along the x-axis. Results shown are from a model with k=12 genetic clusters.

**Figure 3:**
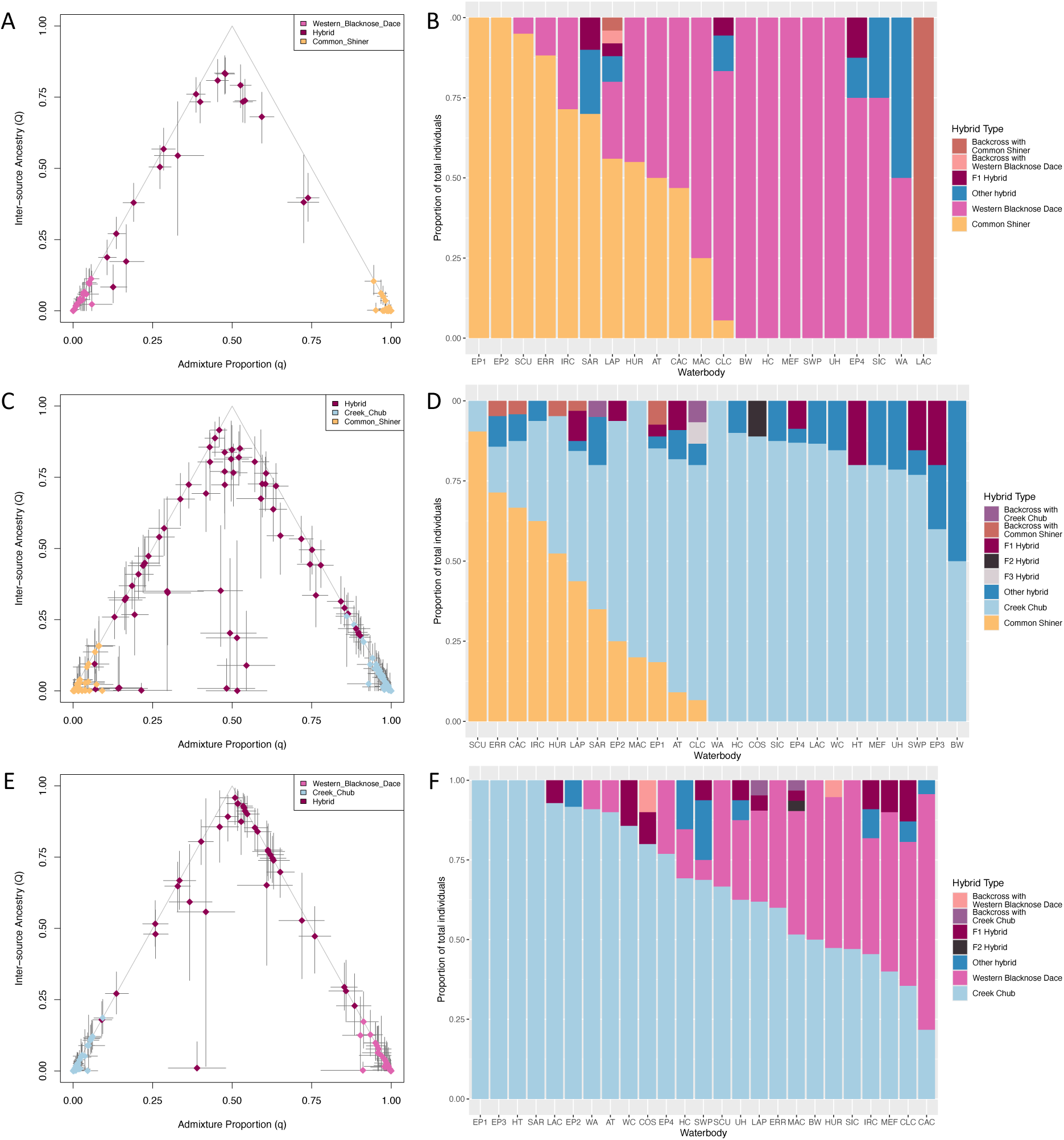
Breakdown of hybridization outcomes for pairwise combinations of common shiner, western blacknose dace, and creek chub. Individuals in the triangle plots are coloured based on their genomic identity from the initial K=12 entropy run. A & B. Western blacknose dace and common shiner. n = 351 individuals total, 30 of which are hybrids. Hybrids and/or parentals were found at 21 of 25 sites. C & D. Creek chub and common shiner. n = 220 individuals total, 18 of which are hybrids. Hybrids and/or parentals were found at all 25 sampling sites. E & F. Western blacknose dace and creek chub. n = 378 individuals total, 49 of which are hybrids. Hybrids and/or parentals were found at all 25 sampling sites.

### 2.5 Assigning genomic identity to individuals

We used custom R scripts to assign a genomic species identification to each individual. This was done using the values of q from entropy, for the best K value, which in this case was K=12. We first extracted the q value data from the HDF5 output files with the R package rhdf5 (Fischer *et al*. 2021, R Core Team 2021) and took the mean of the distribution of q values for each individual.

We then used the q value data to map the species clusters from ENTROPY to phenotypic species, for each value of K. Because phenotypic identifications were done by highly skilled and trained personnel, we assumed that we had correctly identified the majority of individuals using phenotypic traits, therefore, the species with the highest mean proportion of ancestry for a given cluster should be the species which that ancestry is from (Fig. 12SS).

We then assigned each individual a genomic identity based off of admixture proportion from each cluster. If an individual had 90% or greater ancestry from one species, its genomic identity was a parental of that species. This cut-off value has been used in previous studies on hybridization (Hasselman *et al*. 2014), but is more generous than some other studies, which have stricter bounds for classifying parentals (Mandeville *et al*. 2017, Natola *et al*. 2022). If an individual had 10-90% ancestry from exactly two species, it was considered a two-species hybrid between those two species and labeled as a cross between them. If an individual had 10-90% ancestry from more than two species, it was considered a multi-species hybrid and as cross with each parental. Finally, some individuals had 10-90% of only one species; they had 80-90% ancestry from a single source and multiple other smaller contributions from other ancestry sources, precluding them from being either a two-species hybrid or a parental. These individuals were named for their largest ancestry source, but classified as uncertain hybrids, as they were likely the results of many backcrosses with one single species but had ancestors that were different types of F_1_ hybrids and may be effectively acting as parentals. We created a summary of how all individuals were classified in RStudio (R Core Team 2021) with ggplot (Wickham 2016) and dplyr (Wickham *et al*. 2023), and coloured with RColorBrewer (Neuwirth 2014).

Finally, we used the values of q (admixture proportion) and Q (inter-source ancestry) generated by pairwise ENTROPY runs for creek chub, common shiner, and western blacknose dace to classify hybrid individuals by hybrid generation or as backcrosses. To do this, we used a similar custom R script as we used to classify parentals, two-species, and multi-species hybrids. The values for determining which group an individual fell into can be seen in Figure 4S and Table 1S; these values are based on work by Shastry and colleagues (Shastry *et al*. 2021).

**Figure 4:**
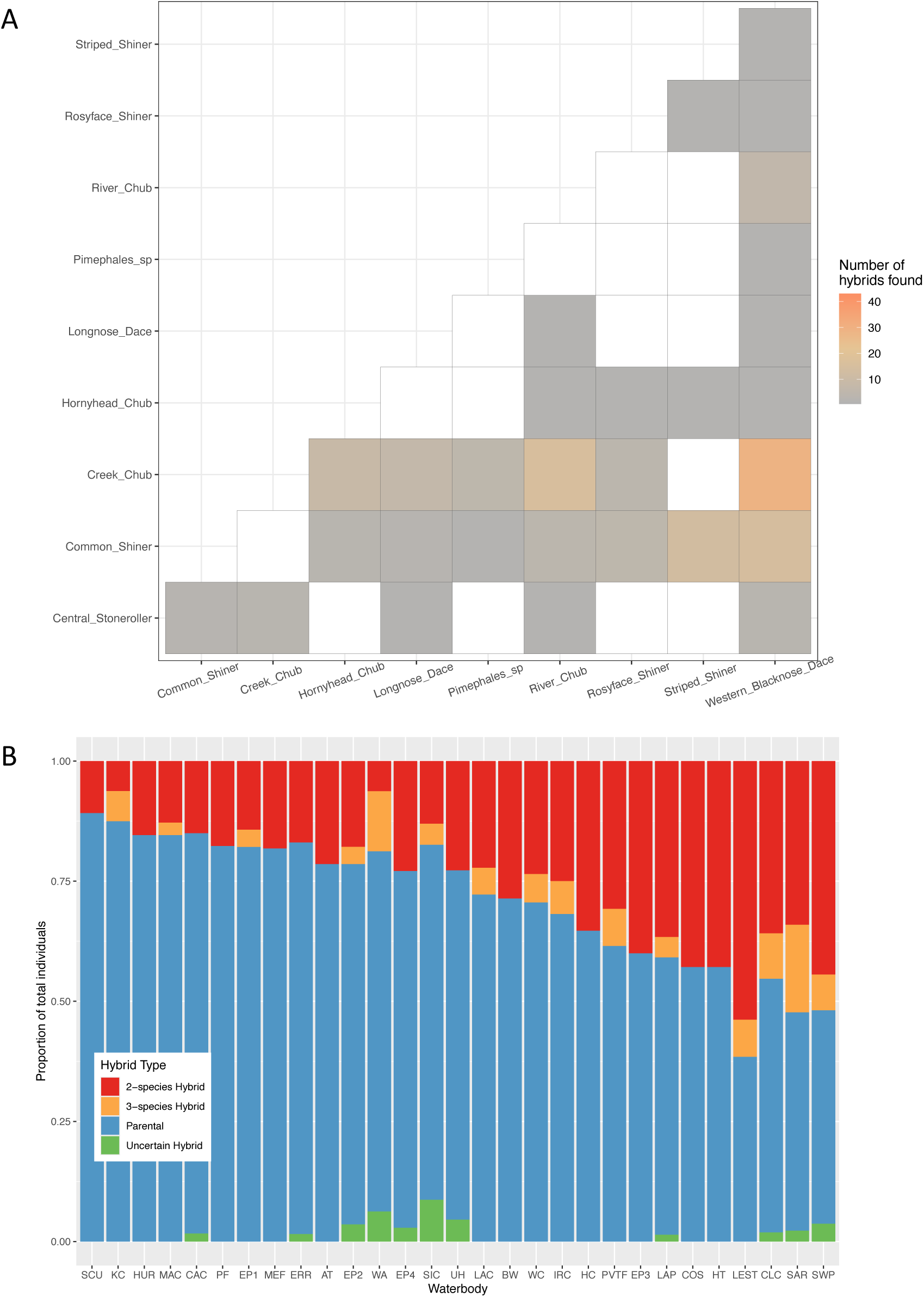
Breakdown of hybridization outcomes for all species crosses and sites. A. Heatmap showing which species that 2-species hybrid crosses were found between and the quantity found. B. Proportion of parental and hybrid individuals, per sampling site.

### 2.6 DNA barcoding

There were eleven individuals identified by ENTROPY to have small proportions of unknown ancestry. We used DNA barcoding in an attempt to assess mitochondrial origin and maternal ancestry and to potentially indicate what species the unanticipated ancestry is from. DNA barcoding was performed by the University of Guelph Advanced Analysis Center Genomics Facility on DNA extracted in the same manner as described above. Both forward and reverse strands of the COI-3 region of the mitochondrial genome was amplified and sequenced using thermalcycler conditions and primers from Ivanova and colleagues (Ivanova *et al*. 2007). We checked the forward fasta sequences with BOLD’s Identification Engine (www.barcodinglife.org/index.php/IDS_OpenIdEngine; Ratnasingham & Hebert 2007) to confirm which species the mtDNA belonged to.

### 2.7 Principal components analysis

We used a principal components analysis (PCA) as a complementary approach to ancestry clustering with ENTROPY. PCA is a model-free approach and allows visualization of genetic differentiation between individuals in multiple dimensions, typically highlighting relationships between species and potentially allowing observations of hybrids between species. We created and visualized principal components analysis in R using RStudio (R Core Team 2021, Posit Team 2025) using a custom R script that calculates a genotype covariance matrix, given genotype point estimates for each locus.

### 2.8 Logistic regression with environmental data

We used a a binomial logistic regression to examine how well leuciscid hybrid ancestry can be predicted by environment of the sampling site’s watershed, including percent land coverage and other characteristics, from OWIT. The response variable was the proportion of individuals classified as hybrids at each site, simplified to amalgamate all the hybrids into one group. We started with 33 predictor variables of environmental data from the Ontario Watershed Information Tool. We then created a correlogram to examine which predictor variables were highly correlated (Fig. 13S). In the case of highly two correlated variables, the one that was judged more likely to have a clear mechanistic link to hybridization outcomes was retained. We ultimately remove 10 of the 33 total variables. We plotted the remaining 23 variables, shown in Figure 14S. We used the glm() function in R (R Core Team 2021) to perform the regression and the ggplot2 package (Wickham 2016) to plot the regression.

Trends in the logistic regressions were driven by few outlier sites with vastly different watershed characteristics, so we also grouped the sites loosely into disturbance categories and ran a logistic regression that plots proportion of hybrids against three categories: agriculturalized sites, urbanized sites, and low-disturbance sites. The two sites from Algonquin Provincial Park (SIC and COS) make up the low-disturbance sites as they are the outlier values in the deciduous treed and clear open water variables, while the three sites from the Waterloo, Ontario region (CLC, LAP, LAC) make up the urbanized category as they have the highest percentages of community infrastructure in their watersheds. Thus, we performed an ANOVA, using a chi-squared test (R Core Team 2021), to determine how strongly the disturbance categories explain variance in the proportion of hybrids per site. Finally, we created four plots per model to assess model fit and spread of residuals, using the plot.lm function in R (R Core Team 2021).

### 2.9 Phylogeny with fishtree

We created a phylogeny of the species sampled for this project, plus fathead minnow and bluntnose minnow in the *Pimephales* genus (Fig. 1C). We used the R package fishtree (Chang *et al*. 2019), which pulls phylogenetic data from its pre-assembled online database, the Fish Tree of Life (fishtreeoflife.org). This database was assembled using DNA sequence data from NCBI GenBank.

## 3 Results

### 3.1 Data filtering

We generated 90,283,210 raw reads from sequencing. Mean alignment of reads to the reference genome, across all 1213 individuals and 9 species, was 43.98% and median alignment was 50.94%. The raw VCF file contained 3,738,553 loci, prior to filtering. After filtering, the VCF file contained 1228 loci and 800 individuals, including all 9 targeted species. Mean alignment of reads to the reference genome was 59.30% after filtering, and median alignment was 65.00% (Fig. 3S). For the two-species cross data sets, the BNDxCC file had 351 individuals, the BNDxCS file had 220 individuals, and the CSxCC file had 378 individuals.

### 3.2 Estimates of ancestry using **ENTROPY**

The DIC value was lowest for the K=12 ENTROPY model (Fig. 5S) and therefore it had a ΔDIC value of 0. This marked K=12 the best model and values from this model were used in further analyses. In the K=12 model, each phenotypically identified species had a reasonably high average q value in separate clusters (Fig. 12SK). In a less optimal model, multiple species may have had high q values in the same column (Fig. 12SA-F, for example). In the K=12 model, however, there were 3 clusters in which no species have high q values: 7, 11, and 12. In Figure 2, cluster 11 and cluster 12 ancestry is uncommon and did not make up a significant portion of ancestry in any one individual. As such, they were both coloured black and not shown in the figure legend. Cluster 7 ancestry (shown in grey) can be seen in individuals at multiple sites that also have creek chub, common shiner, and western blacknose dace ancestry. As well, two individuals were identified as parentals with this ancestry, at two different sites (Fig. 2; sites CLC and UH).

**Figure 5:**
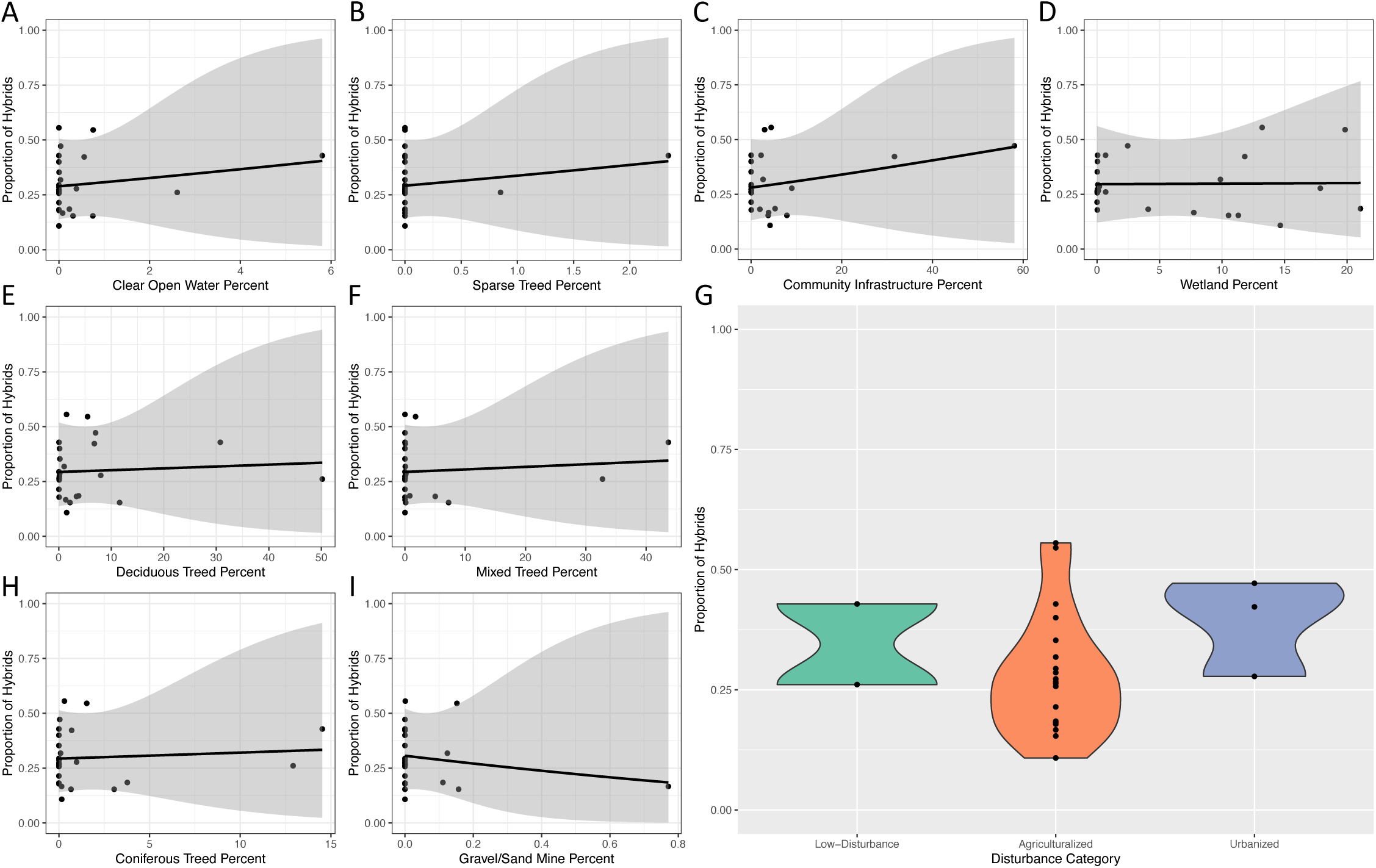
Relationship between proportion of hybrids sampled at a site and land use types. A-F, H, & I. Plots showing a logistic regression of each environmental variable (percent coverage of the watershed with a given land use type) that had a statistically significant relationship with proportion of hybrids per site. n = 25 sites for each plot. A. Clear open water. B. Sparsely covered by trees. C. Community infrastructure. D. Wetlands. E. Deciduous trees. F. Mixed types of trees. H. Coniferous trees. I. Sand and gravel mines or extraction sites. G. Proportion of hybrids sampled at each of the 25 sampling sites, grouped by disturbance type.

By assigning a genomic identity to each individual, we were able to identify the number of two-species hybrids, multi-species hybrids, and parental individuals sampled, as well as deduce the number of individuals that were potentially misidentified in the field (Fig. 15S). Of 731 individuals total, 175 were identified as two-species hybrids (24.0% of individuals) and 40 were identified as multi-species hybrids (5.47% of individuals), for a total of 215 hybrids (29.4% of individuals). Of these 40 multi-species hybrid individuals, 28 were considered proper multi-species hybrids, where more than 2 ancestry sources made up between 10- 90% of these individuals’ ancestry, while 12 had only one ancestry source which contributes between 10-90% of ancestry (Fig. 16S). All “true” multi-species hybrids had 3 main ancestry sources; there were no 4-species or greater hybrids found. All 3-species hybrids matched their phenotypic identity, while 7 of the 12 uncertain hybrids did not (Fig. 15S). The remaining 516 individuals were identified as parental species (70.6% of individuals); of these, 48 of the individuals from parental species did not match their phenotypic identity (9.30% of parentals, Fig. 15S).

We took a mean of the 95% credible intervals (CIs) on each of the 12 q values, for each individual. This resulted in all individuals having one mean CI value. The average mean CI value was 0.00078 and the max was 0.03573 (Fig. 6S). There were 17 individuals that had mean CIs greater than 0.01, most of which (13/17) had at least 10% central stoneroller ancestry. Mean credible intervals for q values in the K=2 species pairs models (Fig. 7S, 8S, 9S) were all much larger than in the K=12 model.

The most commonly sampled two-species hybrid cross was between creek chub and common shiner (Fig. 4A), followed by creek chub and western blacknose dace. Western blacknose dace hybridized with the greatest number of species – all 8 others – followed by common shiner and creek chub which both hybridized with 7 of the 8 other species. Striped shiner hybridized with the fewest other species – only 3 – aside from the *Pimephales* species.

Using ENTROPY, hybrids were identified at every sampling site (Fig. 4B). Most sites (16/25) had both multi-species and two-species hybrids. The proportion of individual fish with hybrid ancestry at a site ranged from 0.108 to 0.556. The sites with the greatest and least proportions of hybrids (SCU and SWP, respectively) were two sites on the same creek. They were originally considered to be one site, but we considered them separate sites for analysis purposes as there was greater than 1km of creek separating them and creek chub rarely disperse further than 1km (Belica & Rahel 2008). Site WA was the only site with a greater proportion of multi-species hybrid individuals than two-species, while site SIC had the same proportions of both types of hybrids.

#### 3.2.1 K=2 comparisons of most abundant species

When we compared our three most common species using pariwise comparisons, we found that creek chub were extremely likely to hybridize with other common species, while common shiner and blacknose dace made fewer hybrids. Specifically, the cross between the two most abundant species, creek chub and common shiner (CCxCS), had both the greatest number of hybrids (49/378) and the greatest proportion of hybrids versus parentals (13.0%) (Table 2S). The western blacknose dace and common shiner cross had the fewest hybrids – only 18/220 (8.2%, Table 2S). Triangle plots displaying these species-pairs analysis can be seen in Figure 3 A,C,E. For BNDxCS, average mean CIs were 0.013 for the value of q and were 0.023 for the value of Q, while maximum CIs were 0.131 for q and 0.270 for Q (Fig. 7S). For CSxCC, average mean CIs for q were 0.025 and for Q were 0.045, while maximum CIs were 0.196 for q and 0.536 for Q (Fig. 8S). The western blacknose dace and creek chub pairing yielded an intermediate number of hybrids – 30/351 individuals or 8.5% – similar to the proportion of western blacknose dace and common shiner hybrids. Average mean Cis for western blacknose dace and creek chub were 0.018 for q and 0.034 for Q, while maximum mean CIs were 0.195 and 0.873, for q and Q, respectively (Fig. 9S). The breakdown of proportion of types of hybrids per site can be seen in Figures Figure 3 B,D, F. While creek chub and western blacknose dace were found at most sites together, common shiner could not always be found with these two species. Despite this, there were still hybrids with common shiner found at most sites.

### 3.3 Principal components analyses

The first two principal components explained 40.395% and 26.81% of variation among the individuals (Fig. 17S A and B). Rosyface shiner, hornyhead chub, and river chub were strongly differentiated from the main cluster of individuals, indicating these groups possessed the greatest amount of differentiation from one another and the rest of the species. In panels C, D, E, and F, PC2 and 3 and PC3 and 4, respectively, central stone roller clustered separately. Finally, in panels G, H, I, and J, the large clump containing creek chub, common shiner, and western blacknose dace began to cluster, as PC5 differentiated these 3 groups despite only accounting for 3.185% of variation. The common shiner, creek chub, and western blacknose dace seemed to form a gradient, in which creek chub laid in the center, but there was some mixing between all three species. Longnose dace separated from the rest of the species very clearly along PC6 (Fig. 17S I, J, K and L). Striped shiner clustered with common shiner and longnose dace clusters close to western blacknose dace, in the phenotypically- coloured plots (Fig. 17SG and I). Interestingly, however, striped shiner never formed a discrete cluster, suggesting that all phenotypically identified striped shiners were actually hybrids. Overall, there was substantial overlap between clusters of species, and none of the clusters were entirely distinct from one another. Resolution of species-specific groups was improved in the genomic identity-coloured plots, as most of the noise between clusters was made up of hybrids.

### 3.4 DNA barcoding reveals hybridization with *Pimephales* species

Eleven individuals were found to have ancestry from a cryptic source population (Fig. 2). The results from DNA barcoding are shown in Table 3S. Of the eleven individuals barcoded, two individuals – AMP22 0463 and EGM19 0302 – showed the presence of mtDNA from two *Pimephales* species: fathead minnow and bluntnose minnow.

### 3.5 Logistic regression shows slight relationships between hybridization and environmental variables

Of the 23 environmental variables that went into the logistic regression, 8 had a statistically significant relationship with the proportion of hybrids per site (Fig. 5). These environmental variables were: percent of land in the watershed covered by clear open water, percent of land in the watershed sparsely covered by trees, percent of land in the watershed covered by deciduous trees, percent of land in the watershed covered by mixed types of trees, percent of land in the watershed covered by coniferous trees, percent of land in the watershed covered by sand and gravel mines or extraction sites, percent of land in the watershed covered by community infrastructure, and percent of land in the watershed covered by wetlands. The estimate, standard error, z-values, and p-values can be seen in table 5S.

The ANOVA with sampling sites categorized into 3 disturbance types showed that the disturbance categories explain variance within the response variable at a statistically significant level (p-value = 0.0008878). Plots created to assess residuals show that the model with the 3 disturbance categories fits the data better than the model with environmental data (Fig. 10S and 11S).

## 4 Discussion

Overall, this study identified extensive and previously undescribed genomic evidence of hybridization within leuciscid minnows. Many hybrids were detected among the sampled individuals (Fig. 2 and hybrids were detected at each sampling site (Fig. 4B), despite no individuals being phenotypically identified as hybrids by skilled field personnel. Some individuals were even found to be multi-species hybrids (Fig. 16S), suggesting a particularly complex history of hybridization. Hybrids were found with two additional species that we had not intentionally included in our sampling (Table 3S); these crosses (Creek Chub x Bluntnose Minnow and Common Shiner with Fathead minnow mitochondrial DNA) were previously unrecorded in literature. Finally, relationships between environmental predictor variables and proportion of hybrids per site were either non-significant, weakly positive, or weakly negative (Fig. 5); this complicated signal indicates that hybridization may not be heavily influenced by anthropogenic disturbances.

### 4.1 Hybridization and admixture occur between almost all sampled species

Species within the Cyprinidae family (which Leuciscidae was only recently distinguished from (Stout *et al*. 2016)) hybridize frequently (Scribner *et al*. 2001, Zbinden *et al*. 2023). Results from ENTROPY show extensive hybridization, similar to a recent report in the literature (Zbinden *et al*. 2023), and many hybrids between more than two species (Fig. 2 and 16S).

All nine phenotypically identified species that were sampled hybridized with at least two other species (Fig. 4A). Of all 45 possible two-species hybrid combinations, 29 (64%) were found in our sampling. Western blacknose dace hybridized with the greatest number of species (all eight others, plus *Pimephales* species), while creek chub and common shiner hybridized with the second most (seven others). Striped shiner hybridized with the fewest other species – only common shiner, rosyface shiner, and hornyhead chub. Despite this, they had the greatest proportion of hybrids to parentals, as in fact no parental striped shiners were present in our data set; all striped shiner ancestry appeared in hybrids (Fig. 2 and 17S).

This finding aligns with results from a comparative study of leuciscid hybridization finding that striped shiners hybridized the most, of the North American minnows they investigated (Corush *et al*. 2020). We also found in our data that most frequently, striped shiners hybridized with common shiner. This is logical as they are sister species (Fig. 1C) and species typically hybridize most with groups they are least diverged from (Bolnick & Near 2005, Presgraves 2002). Additionally, rosyface shiner seemed to rarely hybridize, as few hybrids were found in our data set (Fig. 2), and they were one of the most genetically distinct clusters (Fig. 17S). Despite this, when they did hybridize, it wasn’t limited to a single cross; rosyface shiner hybridized with five other species – a greater number than striped shiner and the same as longnose dace. Finally, because many central stoneroller individuals had very wide 95% credible intervals for genomic identity results, these results involving central stoneroller should be interpreted with caution.

We cannot speculate upon the hybridization dynamics of fathead minnow and bluntnose minnow as we did not sample many parental individuals, and we did not intentionally target these two species, but it was very interesting to see unexpected ancestry from this genus within our data set. DNA barcoding confirmed the presence of mitochondrial DNA from these two *Pimephales* species in individuals containing the cryptic ancestry source, which indicates that these species hybridize with common shiner, creek chub and western blacknose dace (Table 3S). This supported results from ENTROPY, showing that at least 10% *Pimephales* species ancestry is found in 12 individuals – the 11 barcoded plus one multi-species hybrid (Fig. 2). Because *Pimephales* mitochondrial DNA was found in some but not all individuals that ENTROPY showed to have this ancestry source, it indicates that *Pimephales* sp. can be either the maternal or paternal parent of hybrid offspring, unlike systems where one species in a cross must be the maternal parent for the hybrid offspring to be viable (Angers & Schlosser 2007). This makes sense given that both fathead and bluntnose minnow are ingroup among all sampled species (Fig. 1C). While hybrids between bluntnose and fathead minnow with one another and additional species have been found (Heithaus & Grame 1997, Muller & Altadena 2000, Zbinden *et al*. 2023), hybrids with common shiner, creek chub, and western blacknose dace have not previously been recorded in published literature, suggesting that hybridization dynamics among leuciscid minnows are more extensive than previously described.

In addition to the *Pimephales* sp. ancestry found, two additional ancestry sources were identified by entropy (Fig. 12S, clusters 11 and 12). Both of these ancestry sources were found in five species (creek chub, common shiner, western blacknose dace, river chub, and longnose dace) and comprised no more than 1.2% ancestry for any one individual, yet the K=12 model was better fitting to the data than the K=11 or 10 models, according to DIC values. These two sources may represent “ghost admixture”, or ancestry from an unsampled source (Beerli 2004). Given that these segments are so small yet significant enough that their presence is the preferred model, they may come from a fairly divergent source.

The proportion of back crosses and later generation hybrids (F_2_s, F_3_s) found in all three species pairings were about equal, but many more F_1_ hybrids were found with the two crosses with creek chub, than with the BNDxCS cross (Fig. 2S). Hybrids from the BNDxCS cross also have depressed q values in the F_1_ region of the triangle plot, and seem to back cross more towards the common shiner parentals (Fig. 3A). The most likely explanation may be that common shiner was simply more abundant than western blacknose dace, as we caught more of the former than the latter (Fig. 1B). All three species in the K=2 models are non-guarder, brood-hiders, so it is less likely to be a difference in behaviour causing greater reproductive isolation.

There was extensive backcrossing between creek chub and common shiner (Fig. 3C). Many hybrids between these species were classified as later generation back crosses as they fall along the edges of the triangle plot, albeit outside of the ranges to be described as a first-generation backcross. The presence of a wide variety of hybrids and advanced backcrosses indicates that heterozygous genotypes and variably intermediate phenotypes between these species may not be harmful to fitness of these hybrids (Thompson *et al*. 2021). This is quite interesting as western blacknose dace and common shiner share a more recent common ancestor than either one does with creek chub (Fig. 1C), and typically, greater levels of hybridization occur between less diverged species (Bolnick & Near 2005). But given that there was the greatest number of hybrids found between the most abundant species and the least hybrids found between the least abundant species (of the three species for which we analyzed Q values), perhaps divergence is not as important of a reproductive barrier as abundance is, unlike what has been shown in *Centrarchidae* fishes (Bolnick & Near 2005). If all of these species are able to breed with one another, the most common species may be the most likely to hybridize. This fits with the idea that the frequency of hybridization in fish species might be a function of their environment (Hubbs 1955).

The presence of many multi-species hybrids is an interesting and unusual feature of these data (Fig. 16S). We found 40 multi-species hybrids (40/731, 5.5% of all individuals), which were sampled across 16 of the 25 sampling sites (64% of sites). All involved either one ancestry source contributing most of the ancestry plus multiple smaller contributions of ancestry (12/40), or three ancestry sources contributing between 10-90% of ancestry (28/40). Hybridization between multiple species is often overlooked in literature and has traditionally been examined by two species at a time. Recent research has found instances of multi-species hybridization in birds (Natola *et al*. 2022, Ottenburghs 2019), invertebrates (Satokangas *et al*. 2023, Dlouhá *et al*. 2010), and fish (Banerjee *et al*. 2023, Zbinden *et al*. 2023), and there are likely many more multi-species hybrid complexes that exist but have not yet been well studied. Whether hybrids of some species are backcrossing with other species or some species are acting as “hub” species and transmitting ancestry between non-hybridizing species is still unknown (Ottenburghs 2019). However, it is an exciting avenue for further research with these fish.

### 4.2 Hybrids present at all sites

Unlike other studies of hybridization in fish (Banerjee *et al*. 2023, Mandeville *et al*. 2017, Zbinden *et al*. 2023), this system had hybrids at all sample sites and a fairly consistent proportion of hybrid individuals at each site (Fig. 4B). This is true even at sites initially targeted for sampling because of their likelihood to to be much less disturbed by human activity, for example, SIC and COS in Algonquin Provincial Park. This is consistent with morphological evidence stating that these species hybridize naturally, without direct intervention or disturbances by human activity (Cooper 1980, Greenfield *et al*. 1973, Ross 1975, Ross & Cavender 1981). Conversely, site MEF was an extremely disturbed agricultural drainage ditch. While these are very often home to a variety of fishes (Ward-Campbell *et al*. 2017) and are in quite a disturbed area being the target for runoff from fields, relatively few hybrids were found at this site (Fig. 4B).

It is important to note that not all sites are independent - some are hydrologically connected. However, the connected water bodies do not necessarily have identical amounts of hybridization. Sites SCU and SWP are a few kilometers apart along the same stream and yet they had the greatest difference in proportion of hybrids per site (Fig. 2). This could mean that variation in hybridization is super local or that hybrids prefer varied local environments along the length of the stream, much like parentals do. Champagne *et al*. (2022) recently showed that local environment appears to be more impactful on metabolic processes for minnows than watershed-scale environment. Site MEF is connected to site SAR, a much larger river, and there are likely fish that travel between the two. MEF has the advantage where larger minnows such as creek chub may be the top predator in that location (Holm *et al*. 2022, Champagne & McCann 2020). And yet, there were far fewer hybrids found in MEF than SAR and a total lack of multi-species hybrids in MEF. This may perhaps be related to creek chub’s omnivorous behaviour and dominance in the food chain. This trend can be seen more broadly when looking at the common shiner and creek chub crosses, where both parentals are not found together at every site, despite there being hybrids found at most (Fig. 3D). This could potentially be due to behaviours such as dispersal trending towards that of one parent over the other, allowing hybrids to travel to and be captured at sites where one parental tends not to be found, or at least were not present at the time of sampling. It is important to note that sampling always represents a snapshot in time, and might not represent the community present at biologically critical times of year - i.e., when spawning occurs. Regardless of what mechanisms drive hybridization outcomes in this system, it is notable that the types and quantities of hybrids caught at each site were quite variable.

Results of this study emphasize how critical it is to study hybridization dynamics across different areas of the distribution of clades, in part because the same phenotypes of hybrids vary in fitness when placed in varied environmental conditions and geological features. Our study found different results from other recent research on hybridization in leuciscid minnows that was done by Zbinden *et al*. (2023) in across the White River Basin (in Missouri and Arkansas, USA). Zbinden *et al*. (2023) found hybrids at just over half of their sites, while we found hybrids at all sites. While Zbinden *et al*. (2023) included five other families of fish beyond *Leuciscidae*, the authors state that the hybrids they found were overwhelmingly from *Leusiscidae*. They also show creek chub clustering alone with redside dace, separate from central stoneroller and bluntnose minnow, while we found hybrids between creek chub and the latter two species (Fig. 4A). This demonstrates the importance of sampling variable geographic sites when trying to understand hybridization dynamics.

### 4.3 Identification of hybrids in this system benefits from genomics

North American minnow species are difficult to identify. Even with assistance from experi- enced icthyological curators, no sampled individuals used for this project were morphologically identified as hybrids. Results from genomic analysis, however, showed that 29.4% of individuals were hybrids that have at least 10% of ancestry from heterospecific sources and that 9.3% of parentals were misidentified. Hybrids can easily go unnoticed when parental species are difficult to differentiate or when multiple species are involved (Dlouh’a *et al*. 2010, Natola *et al*. 2022, Slager *et al*. 2020) At one time, it was thought that finding hybrids with exclusively intermediate traits precluded the presence of back crossing between those species and vice versa (Hubbs 1955, Greenfield *et al*. 1973). However, the idea that F_1_ hybrids may not look intermediate between parental species has already been demonstrated in plants and fish (Thompson *et al*. 2021, Stelkens & Seehausen 2009). Experimental crosses with creek chub, central stoneroller, hornyhead chub, and redside dace (Ross & Cavender 1981) showed that while hybrids are consistently intermediate between parentals with some characteristics, others may look much more like one parental than the other. This may also be occurring with additional leuciscid fishes.

Hybridization in this clade is not a recent development (Cooper 1980, Greenfield *et al*. 1973, Hubbs 1955, Ross 1975, Ross & Cavender 1981), unlike systems where anthropogenic or other disturbances have altered reproductive isolation and incited hybridization where none existed before (Banerjee *et al*. 2023, Hasselman *et al*. 2014, Natola *et al*. 2022). As such, backcrossing has potentially allowed for a lot of introgression between species. And yet we have defined these fish into species morphologically and used genomic data to show them having separate clusters of ancestry. Thus, there is clearly enough differentiation and reproductive isolation such that genetic drift and selection have formed these fishes into groups with enough genetic differentiation that we describe them as separate species, but not prevented high levels of gene flow. Minnow populations in North America rapidly expanded after the last glacial maxima around 18 thousand years ago (Berendzen *et al*. 2008, Schönhuth *et al*. 2016), and colonization of northern habitat promoted adaptive radiation and speciation events (Bernatchez & Wilson 1998). Understanding the timing of hybridization is critical, because if gene flow between these species has been ongoing and they are so difficult to identify, it begs the question: how can we be sure that hybrids have not influenced our creation of identification tools for these species? F_1_ hybrids are able to be identified in some cases, but backcrosses may look extremely similar to parentals. Traits like lateral scale count are often defined as ranges, and these ranges often overlap (eg. creek chub (52-62), and western blacknose dace (56-68); striped shiner (36-42), common shiner (36-42), and rosyface shiner (37-41) (Holm *et al*. 2022)). Potentially, this could be because hybrid individuals have been unknowingly in the mix when describing morphological traits of these fish. Genomic work can aid in increasing resolution of traits that exist in hybrids compared to parentals and potentially improve the ability to identify and distinguish the two.

### 4.4 Relationship between hybridization and environment remains unclear

We found weak but significant relationships between the environmental data from the OWIT and the proportion of hybrids per site. Most of these relationships were driven by one or two outlier values as there was very little variation within the predictor variables (Fig. 5). These outlier points were typically either sites in Algonquin Provincial Park or the Waterloo area. Because of the clear differences across multiple environmental variables and the lack of strong relationships between proportion of hybrids per site and the environmental variables themselves, we plotted and ran an ANOVA on the proportion of hybrids per site against the sites grouped into three categories, to search for overall trends (Fig. 5G). While there was a significant difference between the mean of the proportion of individuals in the agriculturalized sites and the urbanized sites, the agriculturalized sites had a lower mean proportion of hybrids per site. This was somewhat unexpected and contrasts with our prediction that a greater amount of hybrids would be found at more disturbed sites. However, due to constraints on sampling, the number of sites was not equally distributed across the general disturbance types, hence why we initially performed a logistic regression against the continuous variables from each site, rather than binning them. Binning the sites into categories also reduces the capacity to understand the variation between sites of the same category, which is necessary for the highly agriculturalized region of Southern Ontario, where most watersheds are affected to some degree by agricultural practices.

While it was valuable to explore the use of OWIT data as predictors of hybridization, we ultimately did not see clear, strong results that displayed the extent of the influence of anthropogenic disturbance on hybridization. Sampling an increased number of sites in a greater variety of locations may yield more conclusive results, and indeed similar research in leuciscid minnows in the White River Basin, USA, has shown a positive relationship between precipitation and hybridization (Zbinden *et al*. 2023). While environmental conditions may be different in that area than in Southern Ontario, a connection between the environment and hybridization has been noted in many other fish species, as well (Banerjee *et al*. 2023, Hasselman *et al*. 2014, Seehausen *et al*. 1997). Thus while a strong relationship was not detected here, with increased sampling efforts, we would have more power to test these patterns.

Alternatively, the disturbances that affect hybridization rates in leuciscid minnows in Southern Ontario could be occurring at a much more localized level than included in the data we used. Riparian buffers are important in maintaining stream ecosystem health by filtering runoff and preventing erosion of stream edges (Champagne & McCann 2020, Rahel 2002, Scott & Helfman 2001). Champagne & McCann (2020) found that increased local agricultural activity is more detrimental to terrestrial energy use and trophic position of creek chub than an increase of agricultural activity at the watershed level. In turn, these impacts to metabolic activity in creek chub may also be disturbing their breeding behaviours, in addition to direct disturbance by agricultural practices (Corush *et al*. 2020, Hasselman *et al*. 2014, Seehausen *et al*. 1997). While we did not directly measure riparian zones around the waters we were sampling in this study, comparing riparian zone quality against outcomes of hybridization is an interesting avenue for future work.

Finally, there is also the potential that hybridization this group of fish is simply not affected by anthropogenic disturbances at the levels seen in Southern Ontario. Hybridization produces variable outcomes, especially when looked at across multiple environments with different selective pressures and varying rates of genetic drift and migration (McFarlane *et al*. 2022). Within this noise, anthropogenic disturbance may not be creating clear enough trends to be detectable with the data that we have, especially given how difficult it is to find any truly “undisturbed” sites for comparison with higher disturbance areas. Despite the ultimate limitations on our ability to identify mechanisms controlling hybridization outcomes, we managed to examine hybridization in an extremely complex family across 25 different sites. We have highlighted how variable hybridization outcomes can be across Southern Ontario and have laid the groundwork for future work into mechanisms underlying these hybridization outcomes.

## Acknowledgements

This work was done as a part of the Canada First Research Excellence Program, specifically the University of Guelph’s Food From Thought Program, funded through a Research Support Grant to EGM. The analysis was enabled by a Resources for Research Groups computing al- location to EGM from the Digital Research Alliance of Canada. We would also like to thank the Ministry of Natural Resources and Forestry, Ontario Parks, the Algonquin Wildlife Re- search Station, the Humber River Heritage Trail, the Holland March Drainage System Joint Municipal Service Board, rare Charitable Research Reserve, Pottageville Swamp Conserva- tion area, Scanlon Creek Conservation area, Bruce Municipality, the City of Guelph, and various private land owners, for their assistance with permitting our field work. Finally, we would like to thank members of the McCann and Mandeville labs for their assistance with field work and support during the creation of this manuscript.

## 5 Data Accessibility and Benefit-Sharing

### 5.2 Data Accessibility

Upon the acceptance of this manuscript, data and scripts used for analysis will be made pub- licly available on GitHub (github.com/amanda-meuser) and the NCBI Short Read Archive.

### 5.2 Benefit-Sharing

We collaborated with provincial parks and conservation areas, as well as local property owners, during sampling for this project. Results have been shared with collaborators and the broader scientific community. Genomic data has also been made publicly available and can assist with future work on these species.

## 6 Author contributions

AVM, ARP, and EGM planned and executed sampling strategy. AVM and ARP performed DNA extraction and library preparation. AVM and EGM planned and executed genomic analyses. AVM and SEM modeled environmental correlates of hybridization and statistical analyses. All authors contributed to writing and revising the manuscript.

## Notes

### Competing Interest Statement

The authors have declared no competing interest.

## References

1. Angers B, Schlosser IJ (2007) The origin of Phoxinus eos-neogaeus unisexual hybrids. Molecular Ecology, 16, 4562–4571.

2. Arnold ML, Bouck AC, Cornman RS (2003) Verne Grant and Louisiana Irises: Is there anything new under the sun? New Phytologist, 161, 143–149.

3. Banerjee SM, Powell DL, Moran BM, et al. (2023) Complex hybridization between deeply diverged fish species in a disturbed ecosystem. Evolution, 77, 995–1005.

4. Beerli P (2004) Effect of unsampled populations on the estimation of population sizes and migration rates between sampled populations. Molecular Ecology, 13, 827–836.

5. Belica LA, Rahel FJ (2008) Movements of creek chubs, Semotilus atromaculatus, among habitat patches in a plains stream. Ecology of Freshwater Fish, 17, 258–272.

6. Berendzen PB, Simons AM, Wood RM, Dowling TE, Secor CL (2008) Recovering cryptic diversity and ancient drainage patterns in eastern North America: Historical biogeography of the Notropis rubellus species group (Teleostei: Cypriniformes). Molecular Phylogenetics and Evolution, 46, 721–737.

7. Bernatchez L, Wilson CC (1998) Comparative phylogeography of Nearctic and Palearctic fishes. Molecular Ecology, 7, 431–452.

8. Bolnick DI, Near TJ (2005) Tempo of hybrid inviability in centrarchid fishes (Teleostei: Centrarchidae). Evolution, 59, 1754–1767.

9. Champagne EJ, Guzzo MM, Gutgesell MK, McCann KS (2022) Riparian buffers maintain aquatic trophic structure in agricultural landscapes. Biology Letters, 18.

10. Champagne EJ, McCann KS (2020) *Farming changes the menu for fish: A shift towards autochthonous driven food-webs in agricultural streams*. Ph.D. thesis, University of Guelph, Guelph.

11. Chang J, Rabosky DL, Smith SA, Alfaro ME (2019) An r package and online resource for macroevolutionary studies using the ray-finned fish tree of life. Methods in Ecology and Evolution, 10, 1118–1124.

12. Chunco AJ (2014) Hybridization in a warmer world. Ecology and Evolution, 4, 2019–2031.

13. Cooper JE (1980) Egg, Larval and Juvenile Development of Longnose Dace, Rhinichthys cataractae, and River Chub, Nocomis micropogon, with Notes on Their Hybridization. Copeia, 1980, 469.

14. Corush JB, Fitzpatrick BM, Wolfe EL, Keck BP (2020) Breeding behaviour predicts patterns of natural hybridization in North American minnows (Cyprinidae). Journal of Evolutionary Biology, 34, 486–500.

15. Dlouhá Thielsch A, Kraus RH, Seda J, Schwenk K, Petrusek A (2010) Identifying hybridizing taxa within the Daphnia longispina species complex: A comparison of genetic methods and phenotypic approaches. Hydrobiologia, 643, 107–122.

16. Fischer B, Smith M, Pau G (2021) rhdf5: R Interface to HDF5.

17. Grabenstein KC, Taylor SA (2018) Breaking Barriers: Causes, Consequences, and Experimental Utility of Human-Mediated Hybridization. Trends in Ecology and Evolution, 33, 198–212.

18. Greenfield DW, Abdel-Hameed F, Deckert GD, Flinn RR (1973) Hybridization between Chrosomus erythrogaster and Notropis cornutus (Pisces: Cyprinidae). Copeia, 1, 54–60.

19. Gutgesell MK, Guzzo MM, McCann KS (2025) Agricultural land-use change seasonally rewires stream food webs: a case study from headwater streams in the Lake Erie watershed. Facets, 10.

20. Hasselman DJ, Argo EE, McBride MC, et al. (2014) Human disturbance causes the formation of a hybrid swarm between two naturally sympatric fish species. Molecular Ecology, 23, 1137–1152.

21. Heithaus MR, Grame C (1997) Fish Communities of the Vermilion River Watershed: Comparison of the Main Channel and Tributaries 1. The Ohio Journal of Science, 97, 98–102.

22. Helmers A, Platek A, Ponte M, Secen N, Cottenie K (2016) The impacts of anthropogenic disturbance on plant species richness in the freshwater lakes of Algonquin Provincial Park. SURG Journal, 9, 5–13.

23. Holm E, Burridge ME, Mandrak NE (2022) *A Field Guide to Freshwater Fishes of Ontario*. 3rd edn., Royal Ontario Museum Press.

24. Hubbs CL (1955) Hybridization between Fish Species in Nature. Zoology, 4, 1–20.

25. Ivanova NV, Zemlak TS, Hanner RH, Hebert PD (2007) Universal primer cocktails for fish DNA barcoding. Molecular Ecology Notes, 7, 544–548.

26. Jansson G, Thulin CG, Pehrson A (2007) Factors related to the occurrence of hybrids between brown hares Lepus europaeus and mountain hares L. timidus in Sweden. Ecography, 30, 709–715.

27. Li H (2013) Aligning sequence reads, clone sequences and assembly contigs with BWA-MEM. arXiv, 00, 1–3.

28. Li H, Durbin R (2009) Fast and accurate short read alignment with Burrows-Wheeler transform. Bioinformatics, 25, 1754–1760.

29. Li H, Handsaker B, Wysoker A, et al. (2009) The Sequence Alignment/Map format and SAMtools. Bioinformatics, 25, 2078–2079.

30. Liu WR, Yang YY, Liu YS, et al. (2018) Biocides in the river system of a highly urbanized region: A systematic investigation involving runoff input. Science of the Total Environment, 624, 1023–1030.

31. Mandeville EG, Parchman TL, Thompson KG, et al. (2017) Inconsistent reproductive isolation revealed by interactions between Catostomus fish species. Evolution Letters, 1, 255–268.

32. McFarlane SE, Jahner JP, Lindtke D, Buerkle CA, Mandeville EG (2022) Selection leads to remarkable variability in the outcomes of hybridization across replicate hybrid zones. bioRxiv, pp. 1–18.

33. McIntyre N (2022) *Tourism, Logging, and Community: Finding Balance in South Algonquin and Algonquin Provincial Park* . Ph.D. thesis, Brock University, St. Catherine’s.

34. Meuser AV, Pitura AR, Mandeville EG, Pitura A (2023) A high-quality reference genome for the common creek chub, Semotilus atromaculatus. bioRxiv, pp. 1–23.

35. Moss B (2008) Water pollution by agriculture. Philosophical Transactions of the Royal Society B: Biological Sciences, 363, 659–666.

36. Moyle PB (1999) Effects of Invading Species on Freshwater and Estuarine Ecosystems. In: *Invasive Species and Biodiversity Management*, pp. 177–191, Kluwer Academic Publishers.

37. Muller B, Altadena S (2000) The Mystery of the Feeder Fish, or Who is Rosy Red? American Currents, 26, 19–20.

38. Natola L, Seneviratne SS, Irwin D (2022) Population genomics of an emergent tri-species hybrid zone. Molecular Ecology, 31, 5356–5367.

39. Neuwirth E (2014) RColorBrewer: ColorBrewer Palettes.

40. Ottenburghs J (2019) Multispecies hybridization in birds. Avian Research, 10, 1–11.

41. Parchman TL, Gompert Z, Mudge J, Schilkey FD, Benkman CW, Buerkle CA (2012) Genome-wide association genetics of an adaptive trait in lodgepole pine. Molecular Ecology, **21**, 2991–3005.

42. Posit Team (2025) RStudio: Integrated Development Environment for R.

43. Presgraves DC (2002) Patterns of postzygotic isolation in Lepidoptera. Evolution, 56, 1168– 1183.

44. R Core Team (2021) R: A Language and Environment for Statistical Computing.

45. Rahel FJ (2002) Homogenization of freshwater faunas. Annual Review of Ecology and Sys- tematics, 33, 291–315.

46. Ratnasingham S, Hebert PD (2007) BOLD: The Barcode of Life Data System: Barcoding. Molecular Ecology Notes, **7**, 355–364.

47. Rhymer J, Simberloff D (1996) Extinction by Hybridization and Introgression. Ecology, 27, 83–109.

48. Ross MR (1975) *The Breeding Behaviour and Hybridization Potential of the Norther Creek Chub, Semotilus atromaculatus atromaculatus (Mitchell)*. Ph.D. thesis, Ohio State Uni- versity.

49. Ross MR, Cavender TM (1981) Morphological Analyses of Four Experimental Intergeneric Cyprinid Hybrid Crosses. Copeia, 1981, 377–387.

50. Satokangas I, Nouhaud P, Seifert B, et al. (2023) Semipermeable species boundaries create opportunities for gene flow and adaptive potential. Molecular Ecology, 32, 4329–4347.

51. Schönhuth S, Beachum CE, Knouft JH, Mayden RL (2016) Phylogeny and genetic variation within the widely distributed Bluntnose Minnow, Pimephales notatus (Cyprinidae), in North America. Zootaxa, 4168, 60–38.

52. Schultz K (2003) *Ken Schultz’s Field Guide to Freshwater Fish*. 1st edn., Wiley.

53. Scott MC, Helfman GS (2001) Native Invasions, Homogenization, and the Mismeasure of Integrity of Fish Assemblages. Fisheries, 26, 6–15.

54. Scribner KT, Page KS, Bartron ML (2001) Hybridization in freshwater fishes: a review of case studies and cytonuclear methods of biological inference. Tech. rep.

55. Seehausen O, Van Alphen JJ, Witte F (1997) Cichlid fish diversity threatened by eutrophication that curbs sexual selection. Science, 277, 1808–1811.

56. Shastry V, Adams PE, Lindtke D, et al. (2021) Model-based genotype and ancestry estimation for potential hybrids with mixed-ploidy. Molecular Ecology Resources, 21, 1434–1451.

57. Slager DL, Epperly KL, Ha RR, et al. (2020) Cryptic and extensive hybridization between ancient lineages of American crows. Molecular Ecology, 29, 956–969.

58. Spiegelhalter DJ, Best NG, Carlin BP, Van Der Linde A (2002) Bayesian measures of model complexity and fit. Journal of the Royal Statistical Society Statistical Methodology Series B, 64, 583–639.

59. Stelkens R, Seehausen O (2009) GENETIC DISTANCE BETWEEN SPECIES PREDICTS NOVEL TRAIT EXPRESSION IN THEIR HYBRIDS. Evolution, 63, 884–897.

60. Stout CC, Tan M, Lemmon AR, Lemmon EM, Armbruster JW (2016) Resolving Cyprini- formes relationships using an anchored enrichment approach. BMC Evolutionary Biology, 16, 1–13.

61. Thompson KA, Urquhart-Cronish M, Whitney KD, Rieseberg LH, Schluter D (2021) Pat- terns, Predictors, and Consequences of Dominance in Hybrids. The American Naturalist, 197, 72–88.

62. Todesco M, Pascual MA, Owens GL, et al. (2016) Hybridization and extinction. Evolutionary Applications, 9, 892–908.

63. Wallace AM, Croft-White MV, Moryk J (2013) Are Toronto’s streams sick? A look at the fish and benthic invertebrate communities in the Toronto region in relation to the urban stream syndrome. Environmental Monitoring and Assessment, 185, 7857–7875.

64. Ward-Campbell B, Cottenie K, Mandrak NE, McLaughlin R (2017) Fish assemblages in agricultural drains are resilient to habitat change caused by drain maintenance. Canadian Journal of Fisheries and Aquatic Sciences, 74, 1538–1548.

65. Wickham H (2016) *ggplot2: Elegant Graphics for Data Analysis*. Springer-Verlag New York.

66. Wickham H, Fraņcois R, Henry L, Müller K, Vaughan D (2023) dplyr: A Grammar of Data Manipulation.

67. Wolf DE, Takebayashi N, Rieseberg LH (2001) Predicting the risk of extinction through hybridization. Conservation Biology, 15, 1039–1053.

68. Xu S, Ma J, Ji R, Pan K, Miao AJ (2020) Microplastics in aquatic environments: Occurrence, accumulation, and biological effects. Science of the Total Environment, 703, 1–14.

69. Zbinden ZD, Douglas MR, Chafin TK, Douglas ME (2023) A community genomics approach to natural hybridization. Proceedings of the Royal Society B, 290.

