## Supplemental material for "Extensive multi-species hybridization between Leuciscidae minnow species"

### Supplement

#### 1 PCA with Missing Data

We plotted principal components against each other in a step-wise fashion and coloured them with both phenotypic and genomic identities, to show the contrast between resolution of species clusters. Additionally, to check that principal component value were not simply a reflection of proportion of missing data, we used VCFtools to output a file containing proportion of loci missing data, per individual, and then plotted that against principal components 1 and 2 (Fig.1S).

#### 2 Analysis of breeding behaviour and hybridization

We sorted the individuals into categories of breeding behaviour based on their genomic species identity. For the parentals, we simply placed them into one of three groups based on work done by Corush and colleagues (Corush *et al.* 2020): nest associate (NA), nest builder (NB), or non-guarder, brood-hiders (NGBH). For the hybrids, we categorized them based on the breeding behaviours of their two parental species. We excluded multi-species hybrids from this analysis for sake of simplicity and any parentals or crosses containing *Pimephales* sp. ancestry, as they were not intentionally sampled. The NA group contained only rosyface shiner, while NBs included hornyhead chub and river chub, and finally, NGBHs included all other species: creek chub, common shiner, striped shiner, western blacknose dace, longnose dace, and central stoneroller. Both parentals and hybrids are displayed by number of individuals per group (Fig. 2S). We created the plots in RStudio (R Core Team 2021) using the `R` packages `waffle` (Rudis & Gandy 2019), `hash` (Brown & Hughes 2022), `patchwork` (Pedersen 2023), and `tidyverse` (Wickham *et al.* 2019) and coloured them with `RColorBrewer` (Neuwirth 2014).

We then performed a Pearson's Chi-squared test (Pearson 1900) on the counts of hybrid crosses to see whether there was an excess of any one group that could not be explained by the abundance of parentals of each breeding behaviour. To calculate the expected number of each hybrid cross, we modeled the breeding behaviours as allele frequencies and used a 3-allele version of the Hardy-Weinberg Equilibrium (HWE) equation (Hardy 1908, Mayo 2008, Weinberg 1908):

$$p + q + r = 1$$
$$p^2 + 2pq + 2pr + q^2 + 2qr + r^2 = 1$$

to calculate the "genotype frequencies" – which in this case were the hybrid crosses. We then used the function `chisq.test()` from the base `R` package, `stats` (R Core Team 2021), to compute the Pearson's Chi-squared test.

#### 2.1 Breeding behaviour crosses reflect abundance of parentals

Non-guarder, brood-hider (NGBH) parentals made up the vast majority of individuals sampled (Fig. 2SA). Hybrids between two NGBH individuals were most abundant of all hybrid crosses (Fig. 2SB). The abundance of all other hybrid crosses with NGBHs seemed to follow a similar trend as the abundance of parental types. Nest-builders (NBs) were the second most abundant breeding behaviour in our dataset, and NBxNGBHs were the second most abundant hybrid cross, followed by nest associates (NA) x NGBH. There was only one NAxNB individual and one NBxNB individual. The Pearson's Chi-squared test produced a $\chi^2$ value of 1.8996 and a p-value 0.7542, with 4 degrees of freedom, indicating that the observed counts of hybrid crosses did not deviated from the expected counts in a statistically significant manner.

### References


Brown C, Hughes J (2022) hash: Full Featured Implementation of Hash Tables/Associative Arrays/Dictionaries.

Corush JB, Fitzpatrick BM, Wolfe EL, Keck BP (2020) Breeding behaviour predicts patterns of natural hybridization in North American minnows (Cyprinidae). *Journal of Evolutionary Biology*, **34**, 486–500.

Hardy GH (1908) Mendelian proportions in a mixed population. *Science*, **28**, 49–50.

Mayo O (2008) A century of Hardy-Weinberg equilibrium. *Twin Research and Human Genetics*, **11**, 249–256.

Neuwirth E (2014) RColorBrewer: ColorBrewer Palettes.

Pearson K (1900) On the criterion that a given system of deviations from the probable in the case of a correlated system of variables is such that it can be reasonably supposed to have arisen from random sampling. *The London, Edinburgh, and Dublin Philosophical Magazine and Journal of Science*, **50**, 157–175.

Pedersen TL (2023) patchwork: The Composer of Plots.

R Core Team (2021) R: A Language and Environment for Statistical Computing.

Rudis B, Gandy D (2019) waffle: Create Waffle Chart Visualizations.

Weinberg W (1908) Über den Nachweis der Vererbung beim Menschen. *Jahreshefte des Vereins für vaterländis- che Naturkunde*, **64**, 369–382.

Wickham H, Averick M, Bryan J, *et al.* (2019) Welcome to the Tidyverse. *Journal of Open Source Software*, **4**, 1686.

Table 1S: Parameters for classifying hybrid individuals using both q and Q values, in a K=2 model. Any individual that does not fall within both bounds of either parameter is classified as "Other". BC1P1 = first generation back cross with parental 1, BC1P2 = first generation back cross with parental 2, F1 = first generation hybrid, F2 = second generation hybrid, F3 = third generation hybrid. A visualization of these parameters can be seen in Fig. 4S.

| Hybrid Type | q values | Q values |
|---|---|---|
| Parental 1 | 0 - 0.1 | 0 - 0.2 |
| BC1P1 | 0.1875 - 0.3125 | 0.375 - 0.625 |
| F1 | 0.375 - 0.625 | 0.75 - 1 |
| F2 | 0.375 - 0.625 | 0.375 - 0.625 |
| F3 | 0.375 - 0.625 | 0.25 - 0.375 |
| BC1P2 2 | 0.6875 - 0.8125 | 0.375 - 0.625 |
| Parental 2 | 0.9 - 1 | 0 - 0.2 |

Table 2S: Results from classifying individuals in the K=2 `ENTROPY` models with parameters in Table 1S. BC1P1 = first generation back cross with parental 1, BC1P2 = first generation back cross with parental 2, F1 = first generation hybrid, F2 = second generation hybrid, F3 = third generation hybrid.

| Hybrid Type | Western Blacknose Dace x Creek Chub | Western Blacknose Dace x Common Shiner | Common Shiner x Creek Chub |
|---|---|---|---|
| BC1P1 | 2 | 2 | 2 |
| BC1P2 | 2 | 1 | 6 |
| F1 | 14 | 4 | 12 |
| F2 | 1 | 0 | 1 |
| F3 | 0 | 0 | 1 |
| Other | 11 | 11 | 27 |
| Parental 1 | 96 | 97 | 222 |
| Parental 2 | 225 | 105 | 107 |
| **Total Hybrids:** | 30 | 18 | 49 |
| **Total Parentals:** | 321 | 202 | 378 |

Table 3S: Results from DNA barcoding. A question mark in the third column represents ancestry from an unknown source. Individuals genomically identified as hybrids are shown in the third column, with an "x" between the two greatest sources of ancestry and the source on the left side contributing the greater proportion of ancestry. Two individuals – AMP22_0463 and EGM19_0302 – reveal mtDNA from *Pimephales* species.

| ID Code | Phenotypic Species ID | Genomic Species ID | BOLD DNA Barcode ID |
|---|---|---|---|
| AMP22_0028 | Creek Chub | ? x Creek Chub | Creek Chub |
| AMP22_0463 | Common Shiner | Creek Chub x ? | Bluntnose Minnow |
| AMP22_0511 | Western Blacknose Dace | ? x Western Blacknose Dace | Western Blacknose Dace |
| AMP22_0531 | Western Blacknose Dace | ? | Western Blacknose Dace |
| AMP22_0728 | Common Shiner | Creek Chub x ? | Common Shiner |
| EGM19_0099 | Creek Chub | Creek Chub x ? | Creek Chub |
| EGM19_0302 | Common Shiner | ? | Fathead Minnow |
| EGM19_0393 | Creek Chub | Creek Chub x ? | Creek Chub |
| EGM19_0757 | Creek Chub | ? | Common Shiner |
| EGM19_1049 | Creek Chub | Creek Chub x ? | Creek Chub |
| EGM19_1288 | Common Shiner | ? x Common Shiner | Creek Chub |

Table 4S: Random forest algorithm predictions of proportions of hybrid individuals found at a site, given environmental data from the upstream watershed. Site numbers indicate order in testing data set and are not indicative of waterbody code in any way.

| Site | Actual proportion of hybrid individuals | Predicted proportion of hybrid individuals, pre-optimization | Predicted proportion of hybrid individuals, post-optimization |
|---|---|---|---|
| 1 | 0.214 | 0.290 | 0.276 |
| 2 | 0.400 | 0.284 | 0.281 |
| 3 | 0.154 | 0.326 | 0.275 |
| 4 | 0.108 | 0.259 | 0.288 |
| 5 | 0.556 | 0.259 | 0.288 |

Table 5S: Coefficient scores from the logistic regression, including the estimate, standard error, z-value, and p-value. Significance is denoted with an asterisk (*) if the p-value is less than 0.05, and with two asterisks (**) if the p-value is below 0.01.

| Coefficients: | Estimate | Std. Error | z-value | p-value | Significance Codes |
|---|---|---|---|---|---|
| (Intercept) | -4.975e+02 | 4.417e+02 | -1.126 | 0.26010 | |
| Lat | 1.114e+01 | 8.321e+00 | 1.339 | 0.18063 | |
| Long | -7.162e-02 | 2.484e+00 | -0.029 | 0.97700 | |
| Drainage_area_km2 | -3.311e-03 | 5.151e-03 | -0.643 | 0.52043 | |
| Length_main_channel_km | -2.350e-02 | 1.251e-01 | -0.188 | 0.85097 | |
| Max_channel_elevation_m | -2.005e-02 | 8.206e-02 | -0.244 | 0.80701 | |
| Min_channel_elevation_m | 1.504e-02 | 1.087e-01 | 0.138 | 0.88994 | |
| Mean_elevation_m | -9.192e-02 | 1.190e-01 | -0.773 | 0.43975 | |
| Max_elevation_m | 4.548e-02 | 1.017e-01 | 0.447 | 0.65490 | |
| Mean_slope_percent | 3.186e-01 | 9.522e-01 | 0.335 | 0.73796 | |
| Annual_precipitation_mm | 3.686e-02 | 3.886e-02 | 0.948 | 0.34289 | |
| Clear_Open_Water_percent | 7.680e+00 | 3.345e+00 | 2.296 | 0.02166 | * |
| Sparse_Treed_percent | 6.133e+01 | 3.060e+01 | 2.004 | 0.04507 | * |
| Treed_Upland_percent | 2.742e+01 | 1.761e+01 | 1.557 | 0.11937 | |
| Deciduous_Treed_percent | 1.942e+00 | 9.132e-01 | 2.126 | 0.03349 | * |
| Mixed_Treed_percent | -7.377e+00 | 3.444e+00 | -2.142 | 0.03220 | * |
| Coniferous_Treed_percent | 3.834e+00 | 1.863e+00 | 2.058 | 0.03956 | * |
| Plantations_Treed_Cultivated_percent | -2.139e+00 | 1.430e+00 | -1.496 | 0.13462 | |
| Hedge_Rows_percent | -3.443e+00 | 3.299e+00 | -1.043 | 0.29672 | |
| Tallgrass_Woodland_percent | 7.845e+02 | 1.565e+03 | 0.501 | 0.61612 | |
| Sand_Gravel_Mine_Tailings_Extraction_percent | -1.095e+01 | 4.634e+00 | -2.362 | 0.01819 | * |
| Community_Infrastructure_percent | -2.465e-01 | 9.472e-02 | -2.603 | 0.00925 | ** |
| Agriculture_and_Undifferentiated_Rural_Land_Use_percent | 8.069e-02 | 6.139e-02 | 1.314 | 0.18870 | |
| Total_wetland_percent | -8.358e-01 | 3.515e-01 | -2.378 | 0.01740 | * |

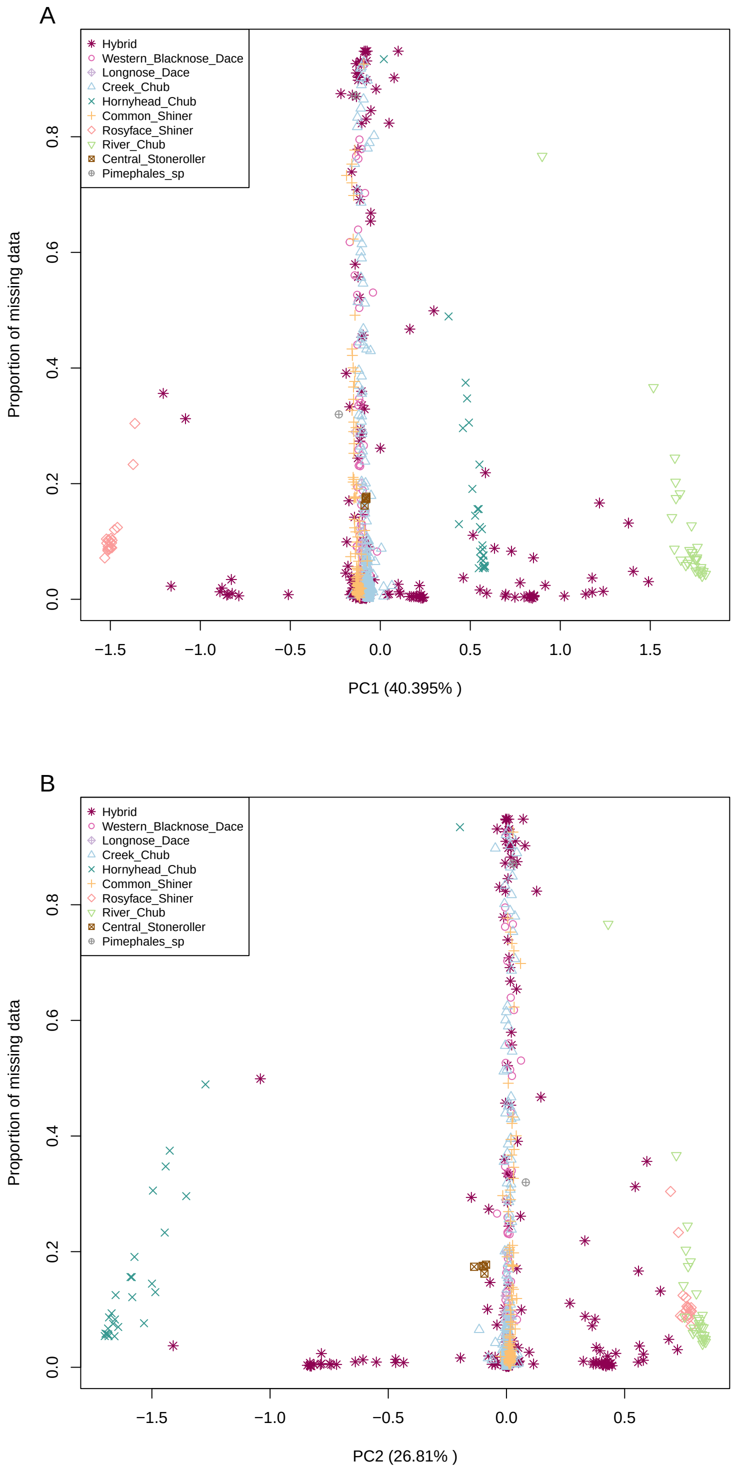


Figure 1S: Proportion of missing data plotted against principal component. Individuals are coloured by genomic identity. A. Principal component 1. B. Principal component 2.

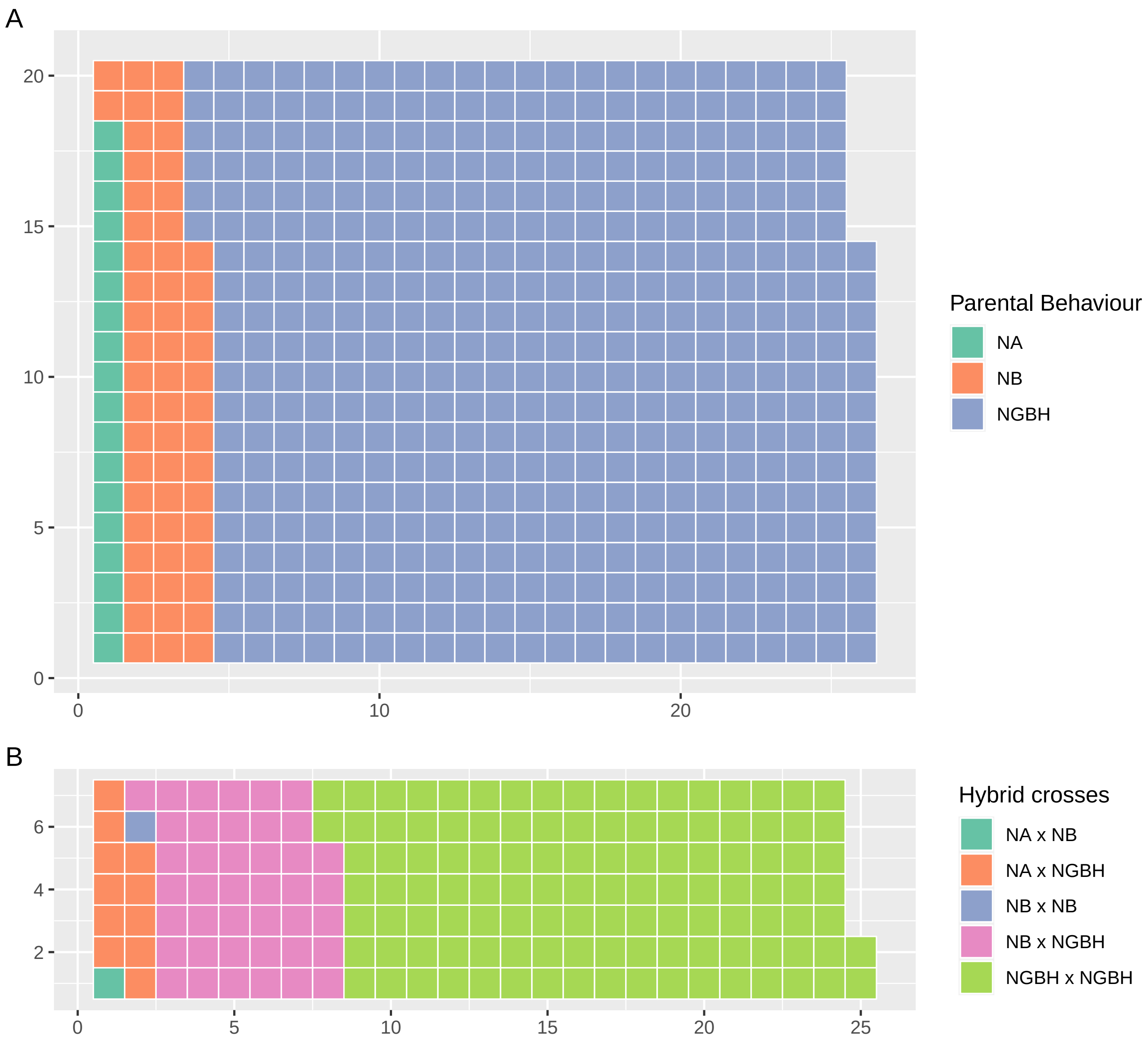


Figure 2S: Waffle plots depicting the number of individuals that display each breeding behaviour or their parentals' breeding behaviours. One square represents one individual. Any parentals or hybrid crosses including *Pimephales* sp. were excluded. NA = nest associate, NB = nest builder, NGBH = non-guarder, brood-hider. A) Parental individuals, n = 514. B) Hybrid crosses, n = 170.

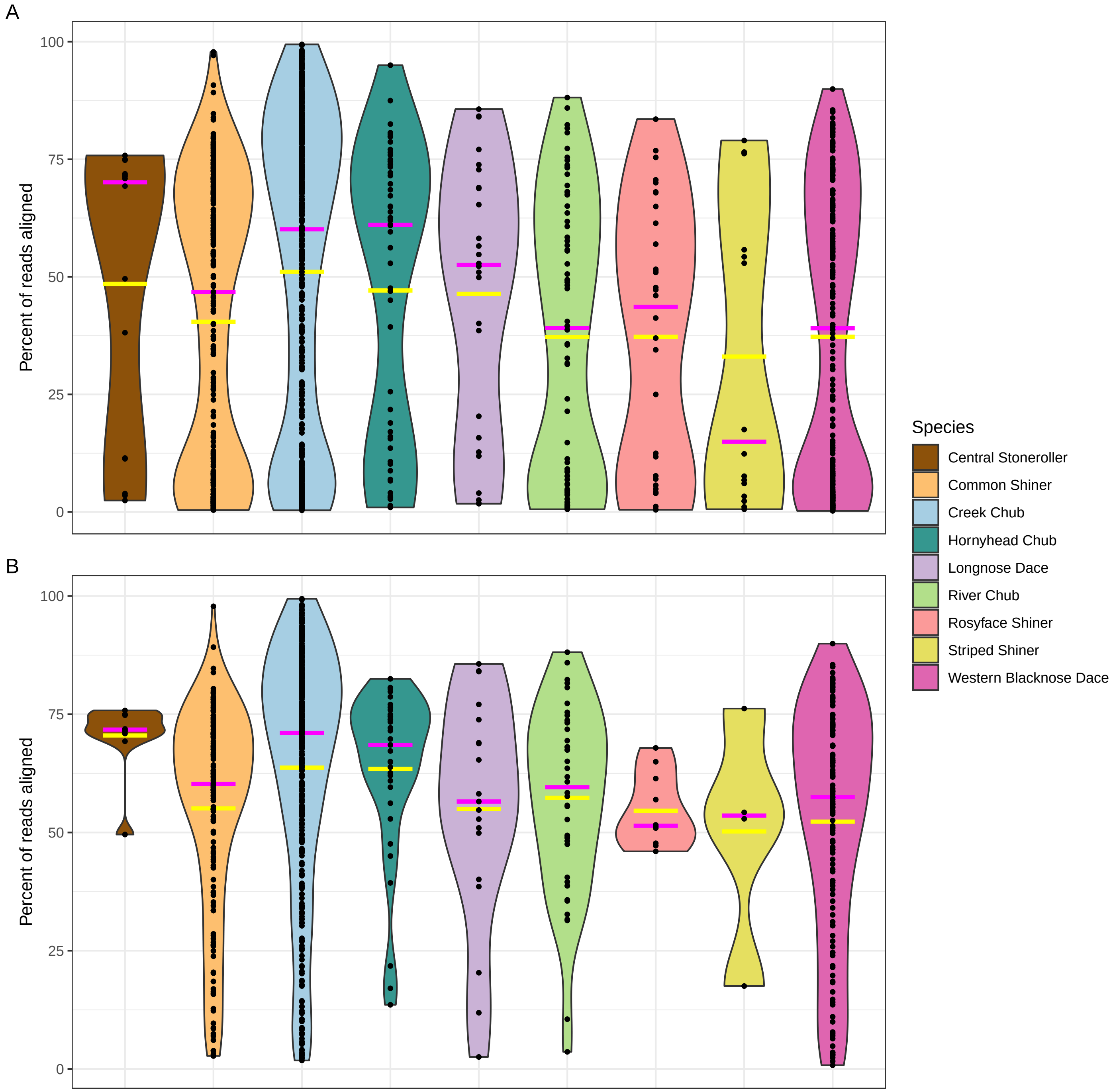


Figure 3S: Violin plots showing rates of alignment to the reference genome, per phenotypically identified species. Yellow bars indicate mean and magenta bars indicate median. A. Prior to filtering, overall mean alignment is 43.98% and median is 50.94%. n = 1213. B. Post-filtering, overall mean alignment is 59.30% and median is 65.00%. n = 731.

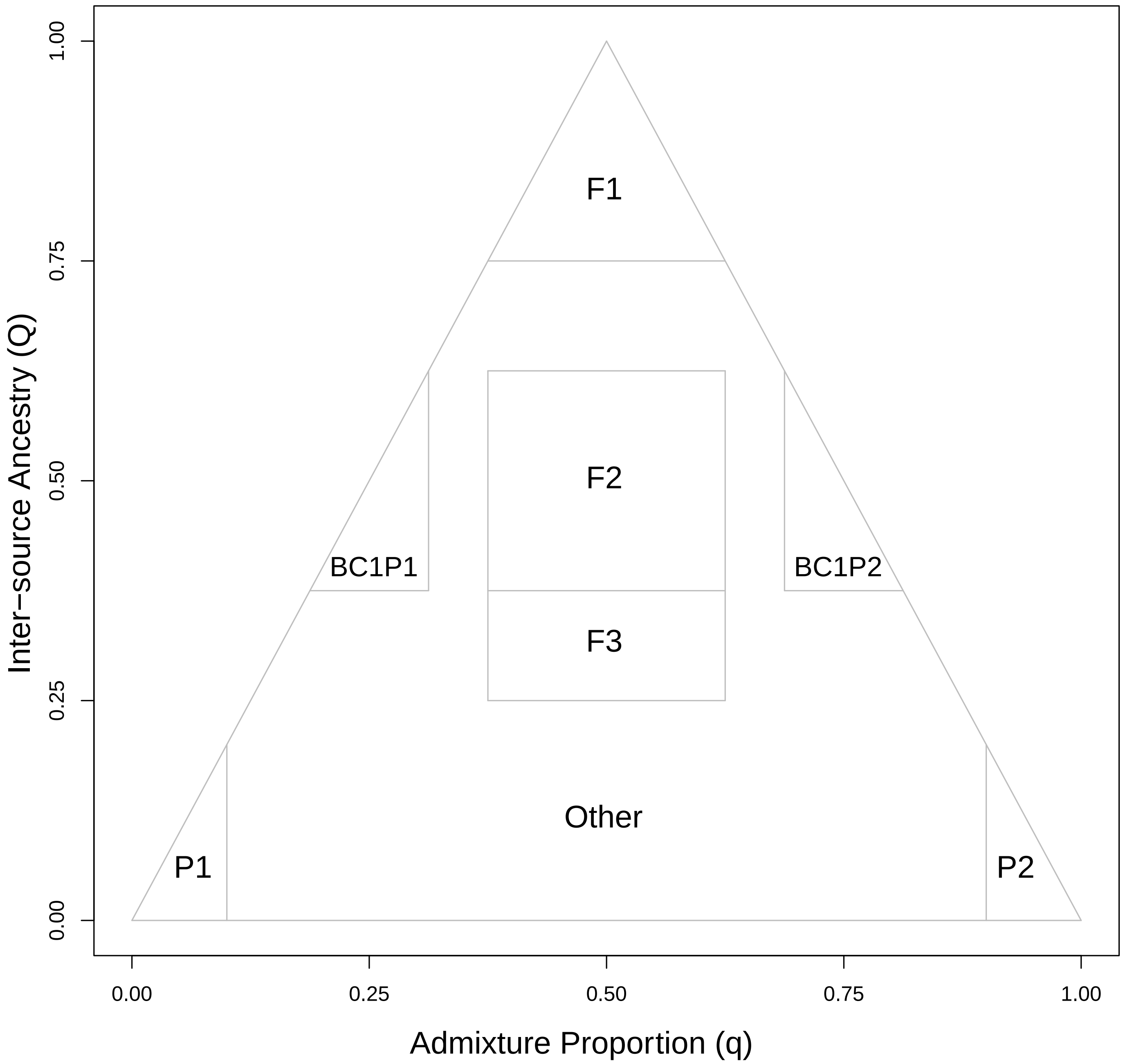


Figure 4S: Parameters for classifying hybrid individuals using both q and Q values, in a K=2 model. Any individual that does not fall within both bounds of a specified region is classified as "Other". BC1P1 = first generation back cross with parental 1, BC1P2 = first generation back cross with parental 2, F1 = first generation hybrid, F2 = second generation hybrid, F3 = third generation hybrid. Values delimiting each region can be found in Table 1S.

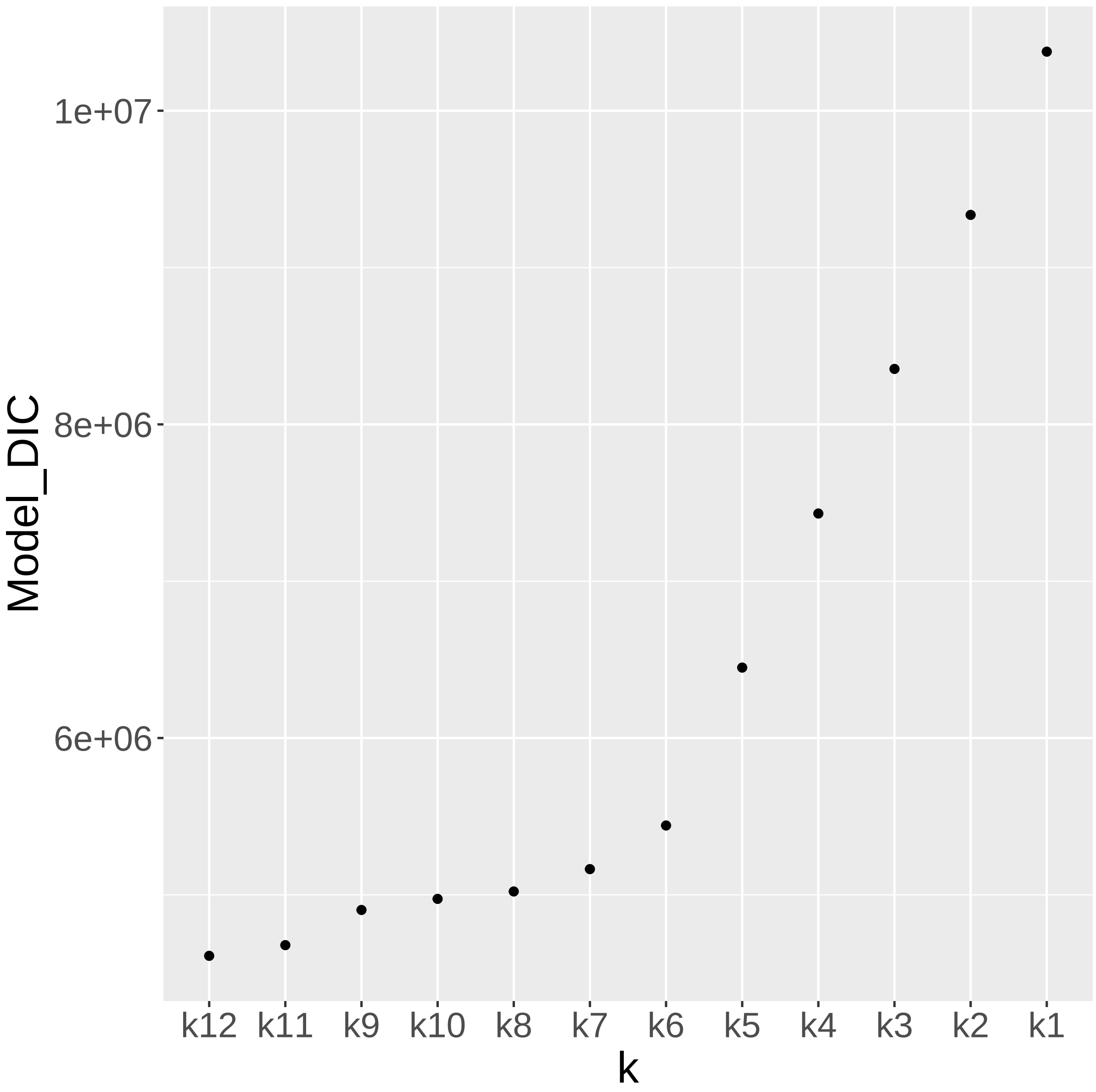


Figure 5S: Deviance information criterion (DIC) plotted for each value of k. K=12 has the lowest value, therefore has a delta DIC value of 0 and is the best value of k.

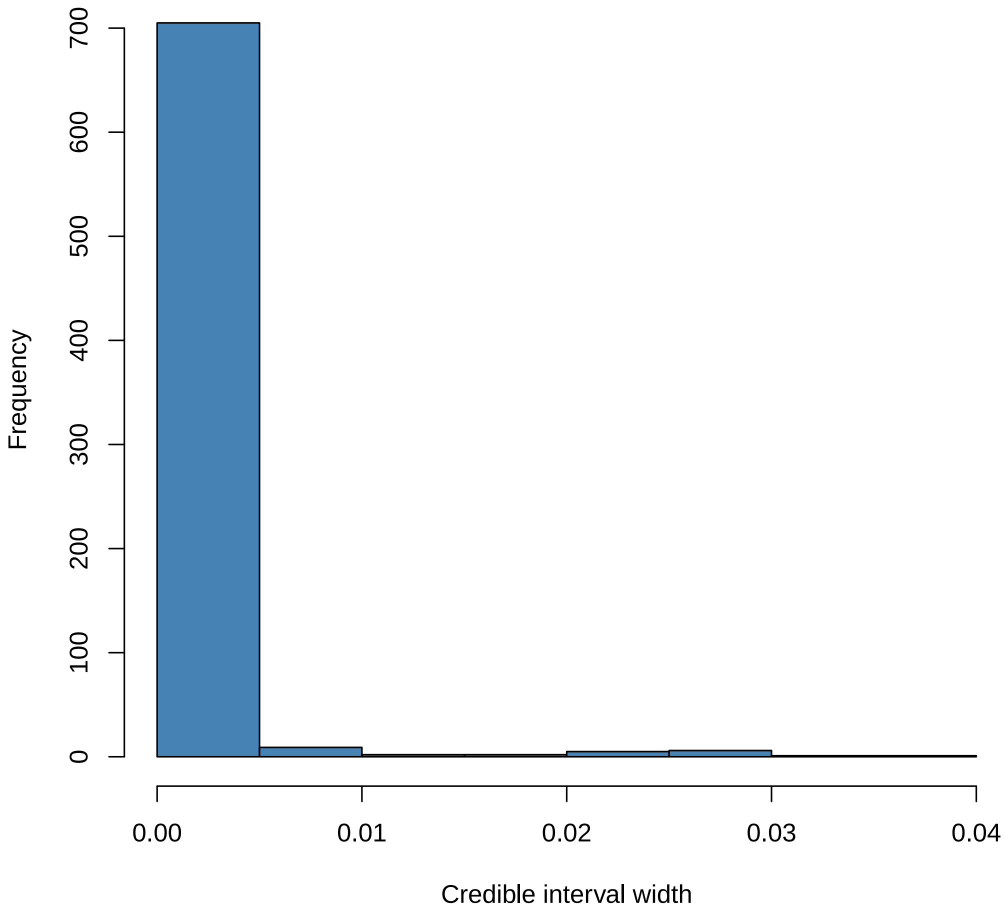


Figure 6S: Histogram showing the mean credible interval width for each individual in the K=12 model. The average mean CI value was 0.00078 and the max was 0.03573.

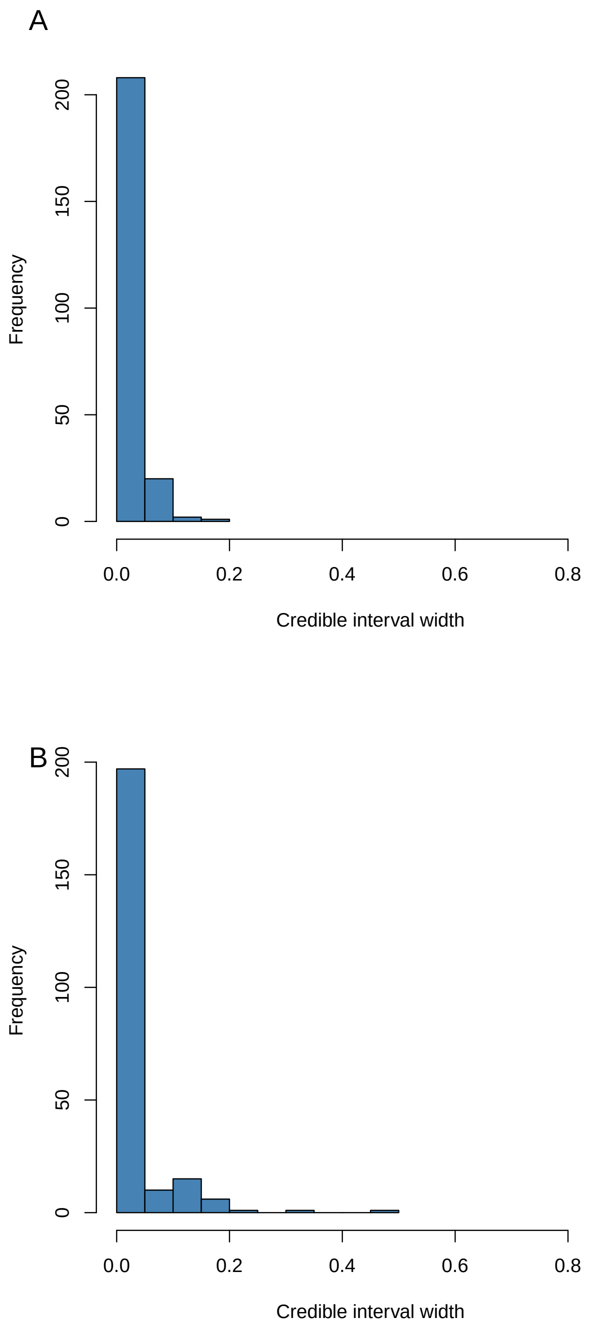


Figure 7S: Histogram showing the mean credible interval (CI) width for each individual in the BNDxCS model. A. Credible intervals for q. The average mean CI value was 0.013 and the max was 0.131. B. Credible intervals for Q. The average mean CI value was 0.023 and the max was 0.270.

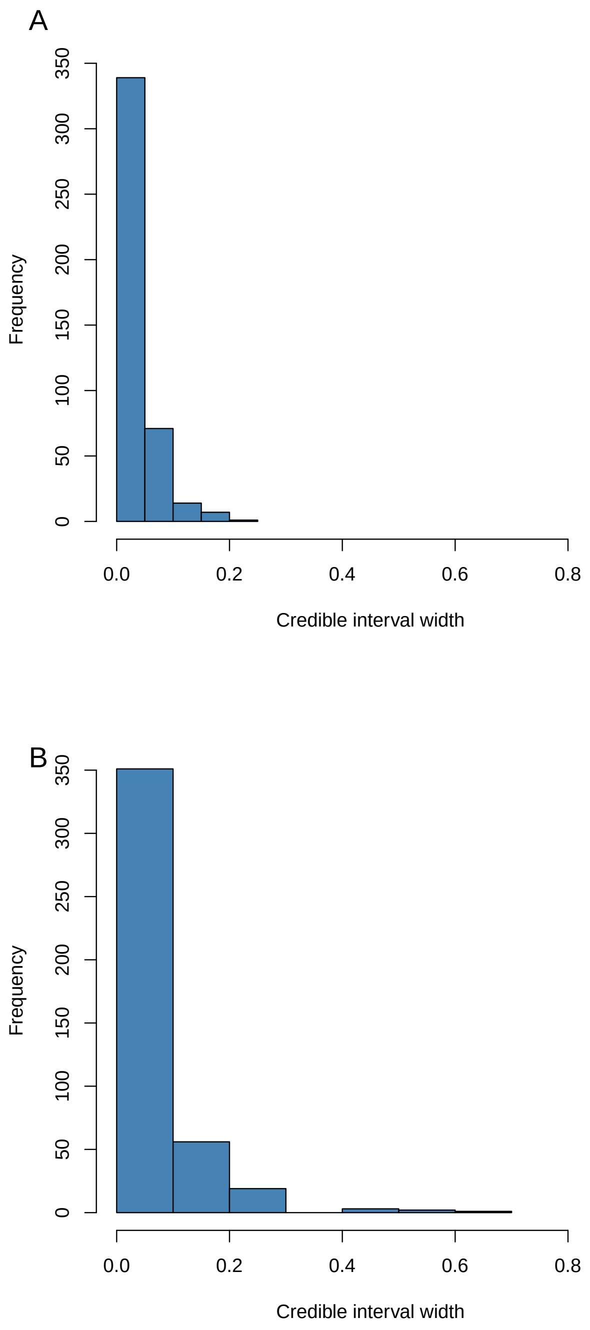


Figure 8S: Histogram showing the mean credible interval (CI) width for each individual in the CSxCC model. A. Credible intervals for q. The average mean CI value was 0.025 and the max was 0.196. B. Credible intervals for Q. The average mean CI value was 0.045 and the max was 0.536.

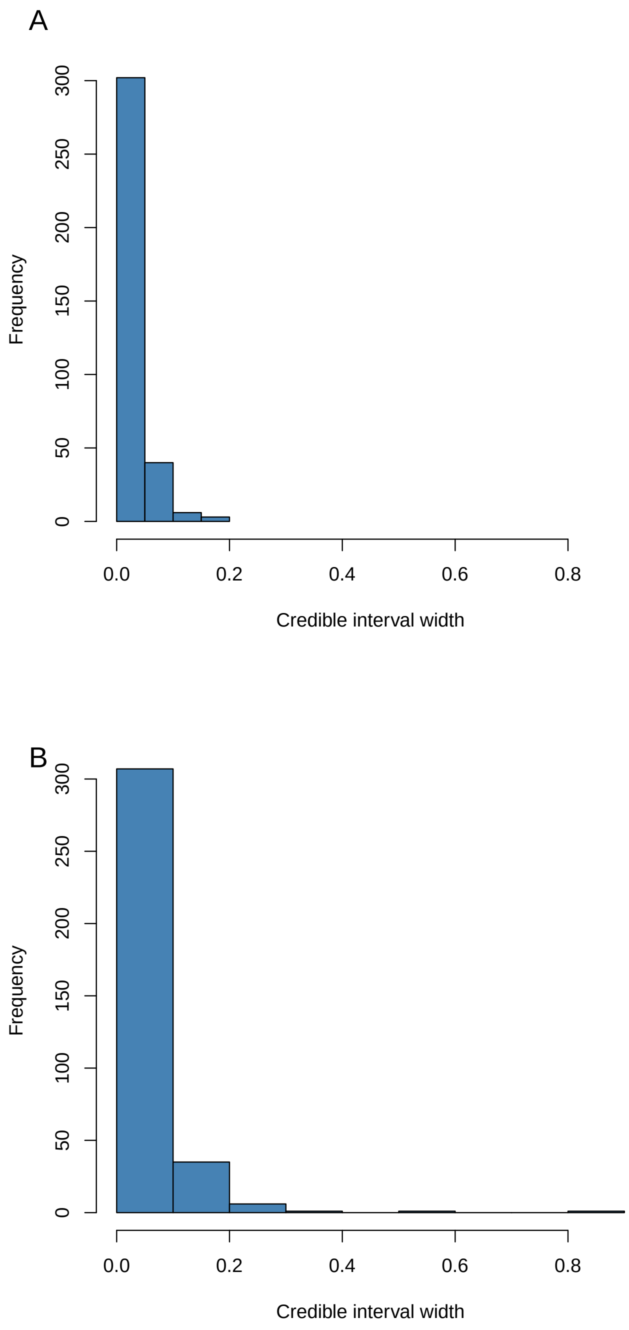


Figure 9S: Histogram showing the mean credible interval (CI) width for each individual in the BNDxCC model. A. Credible intervals for q. The average mean CI value was 0.018 and the max was 0.195. B. Credible intervals for Q. The average mean CI value was 0.034 and the max was 0.873.

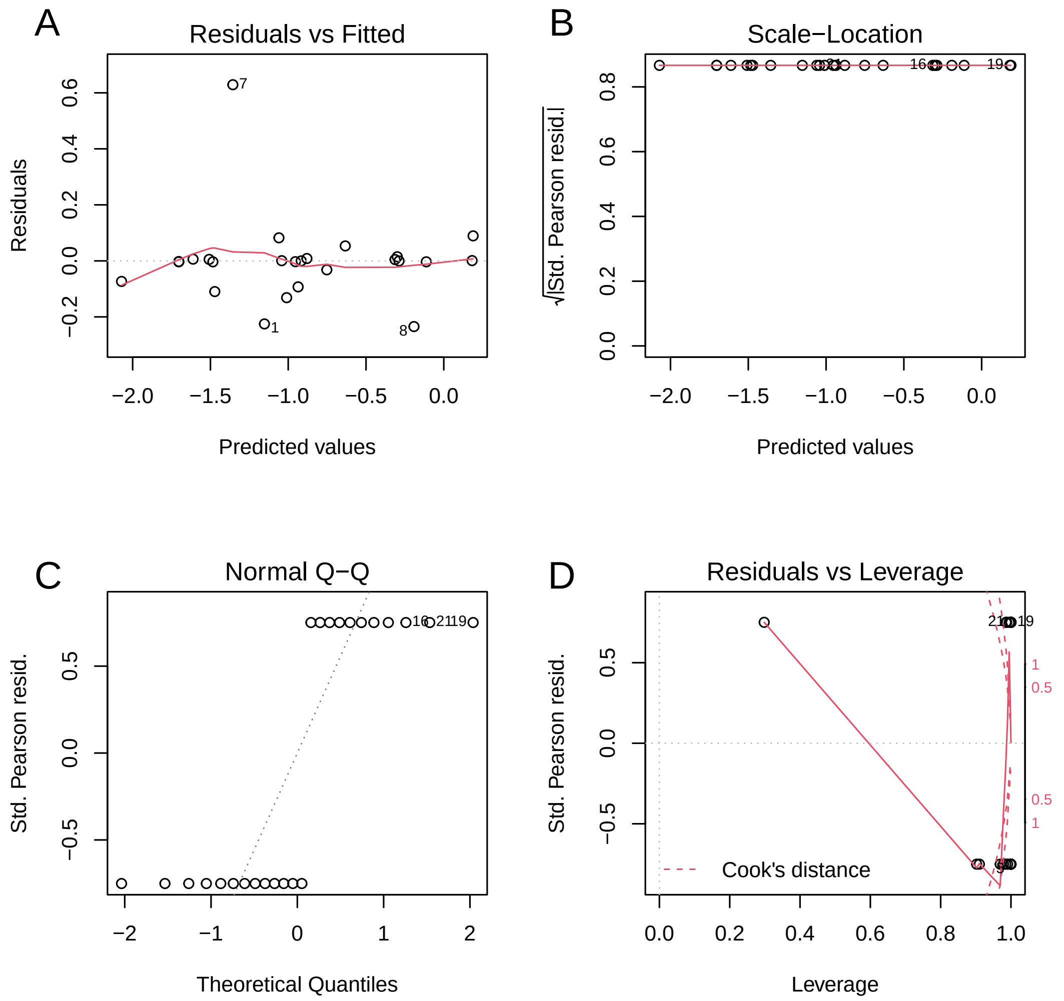


Figure 10S: Plotted residuals for logistic regression with environmental variables and proportion of hybrids at each of the 25 sampling sites.

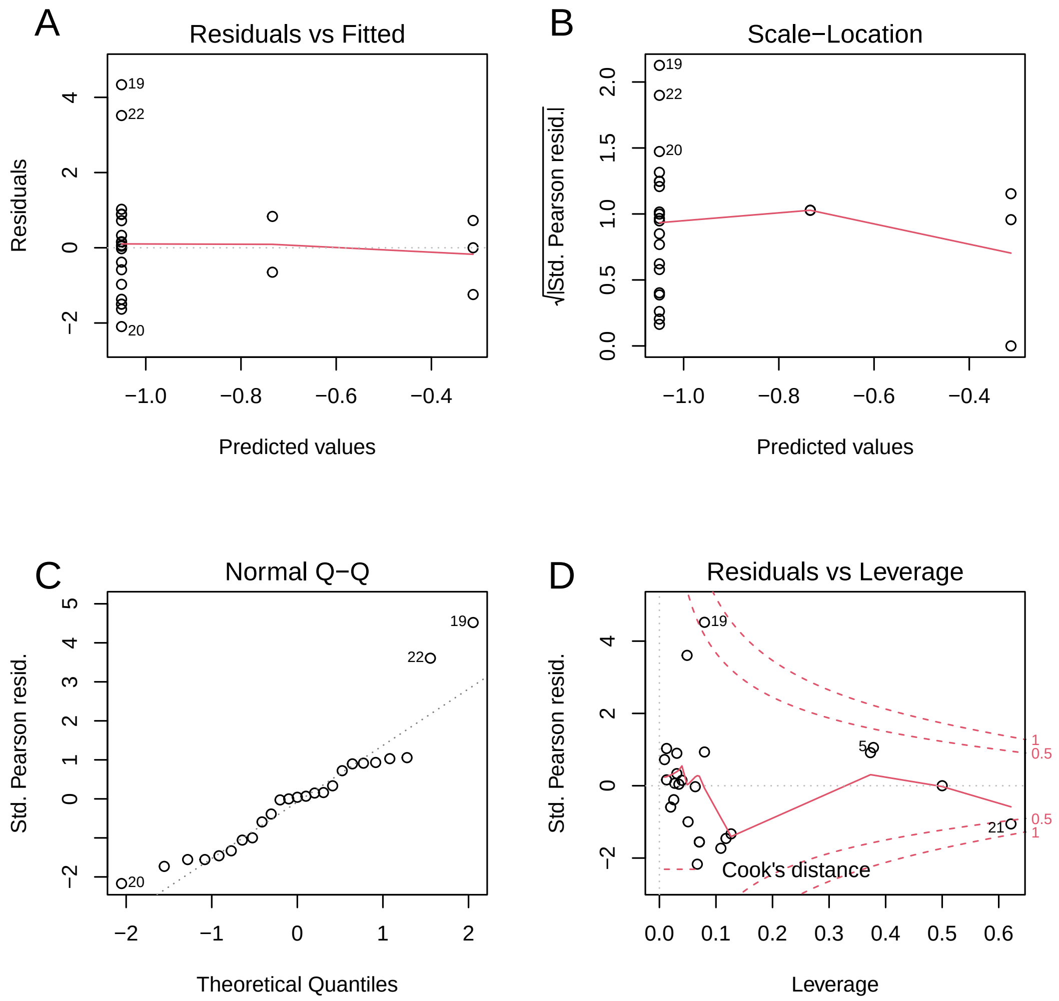


Figure 11S: Plotted residuals for logistic regression with proportion of hybrids at each of the 25 sampling sites, grouped by disturbance type.

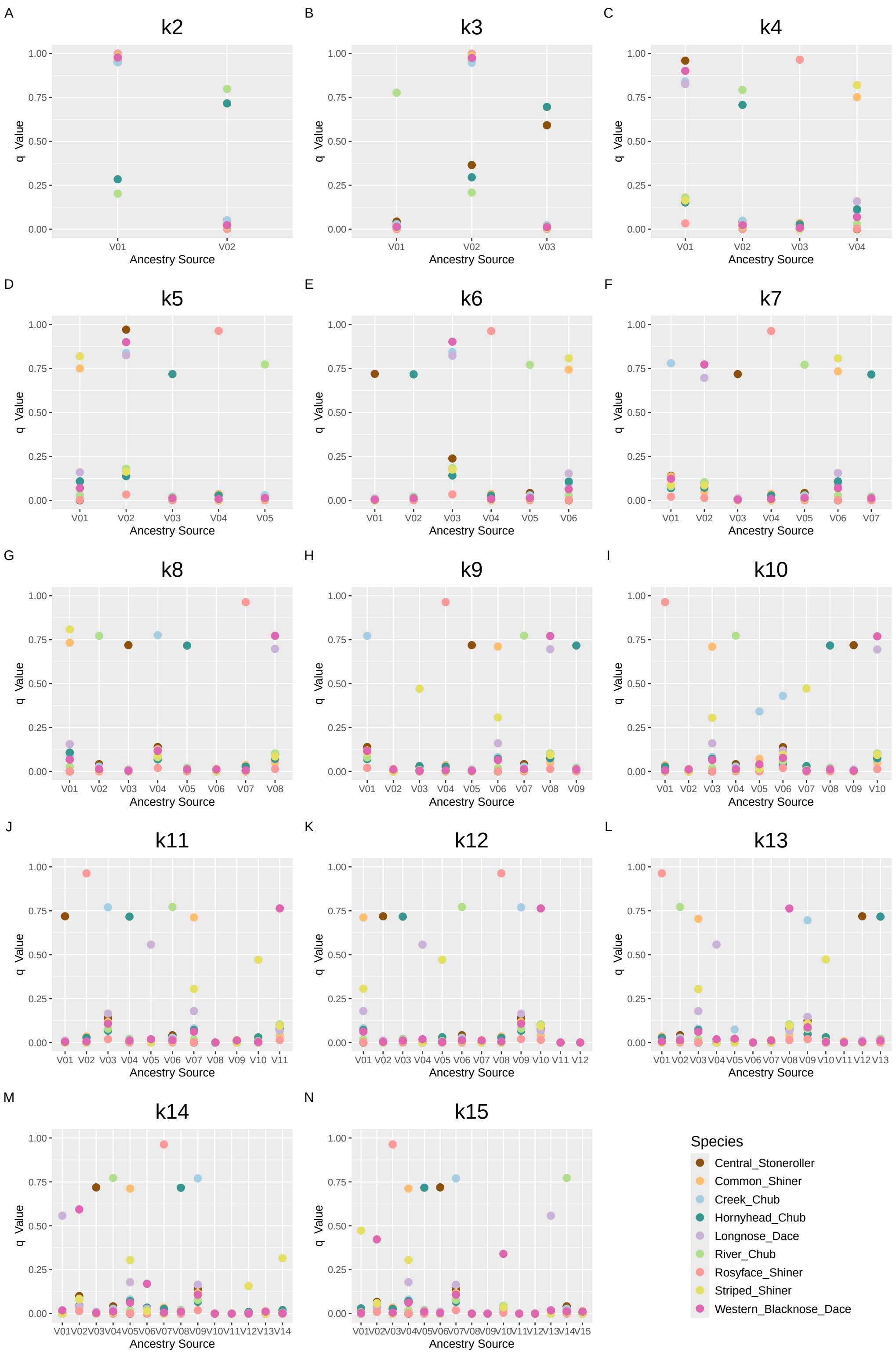


Figure 12S: Mean q value of all individuals in each phenotypically identified species, for all values of K ran through ENTROPY. Subplot K shows the results for K=12, which was the best model.

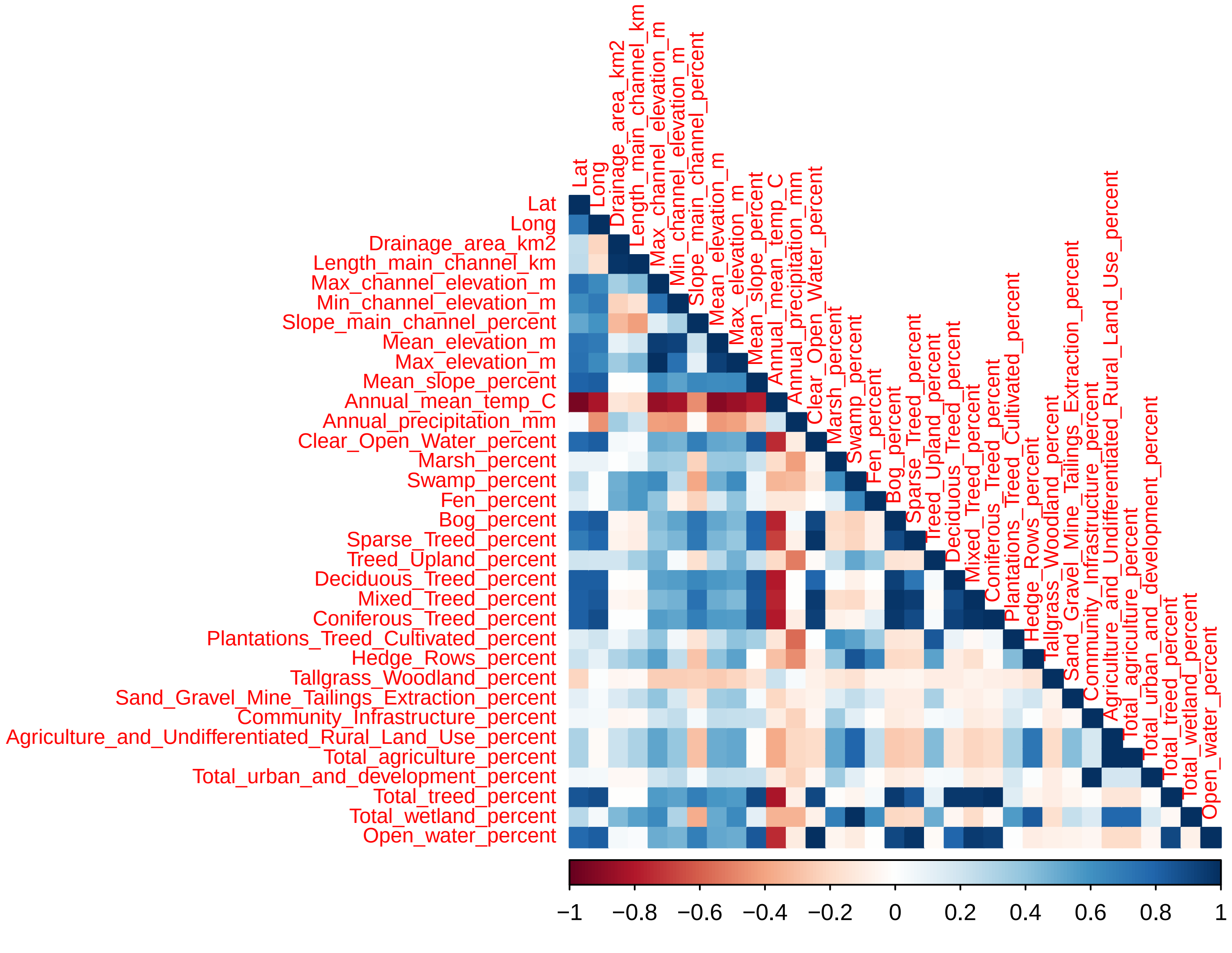


Figure 13S: Correlogram showing correlation between 33 variables from the Ontario Watershed Information Tool. Red colouring indicates a negative correlation between variables while blue colouring indicates a positive relationship.

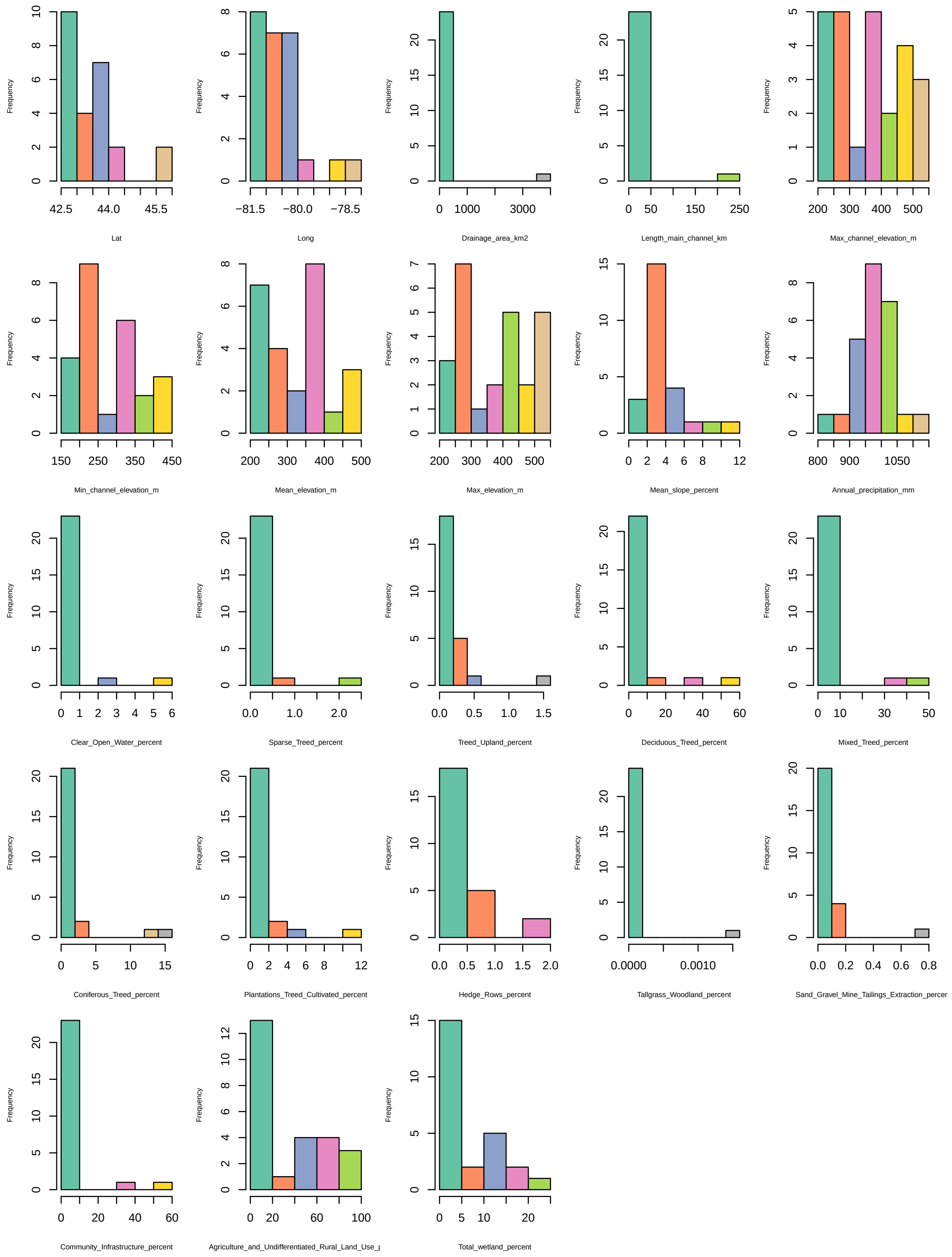


Figure 14S: Histograms showing the 23 variables from the Ontario Watershed Information Tool, used in the logistic regression.

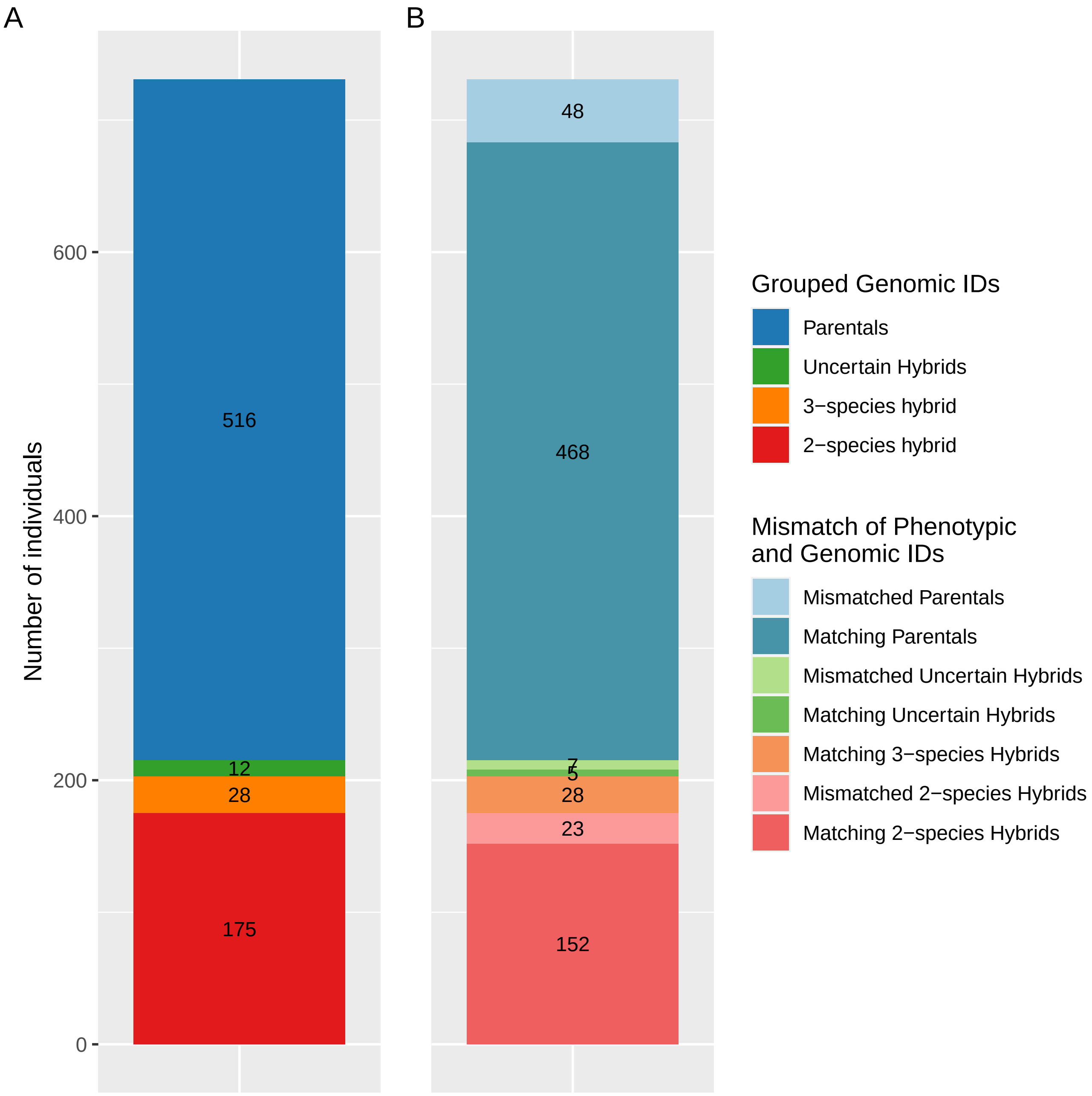


Figure 15S: Summary of genomic identities assigned and number of individuals with discordant genomic and phenotypic identities. A. Summary of genomic identities as determined by `ENTROPY`. B. Comparison of genomic identities to initial phenotypic identities from field work. All hybrids were initially considered misidentified, as no individuals were actually identified as a hybrid during sampling, but were considered to match their phenotypic identity if one of the species in the hybrid cross matched the species of phenotypic identity. Individuals considered "mismatched" are discussed at greater length in Section **??**. n = 731

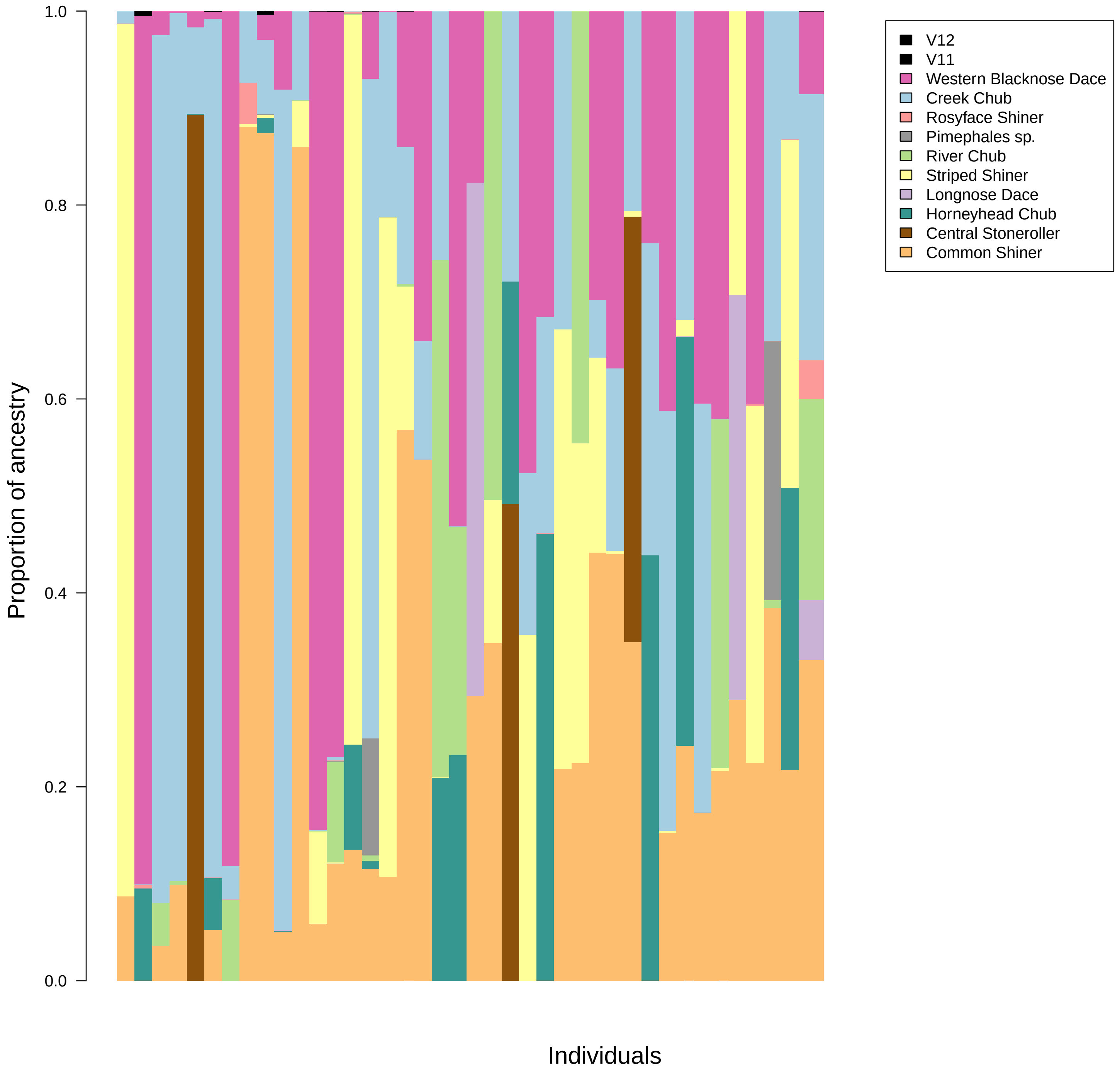


Figure 16S: Bar plot of q values for multi-species hybrids. This includes both the uncertain hybrids and the 3-species hybrids. Individuals are ordered from left to right by the sizes of their largest proportion of ancestry. n = 40

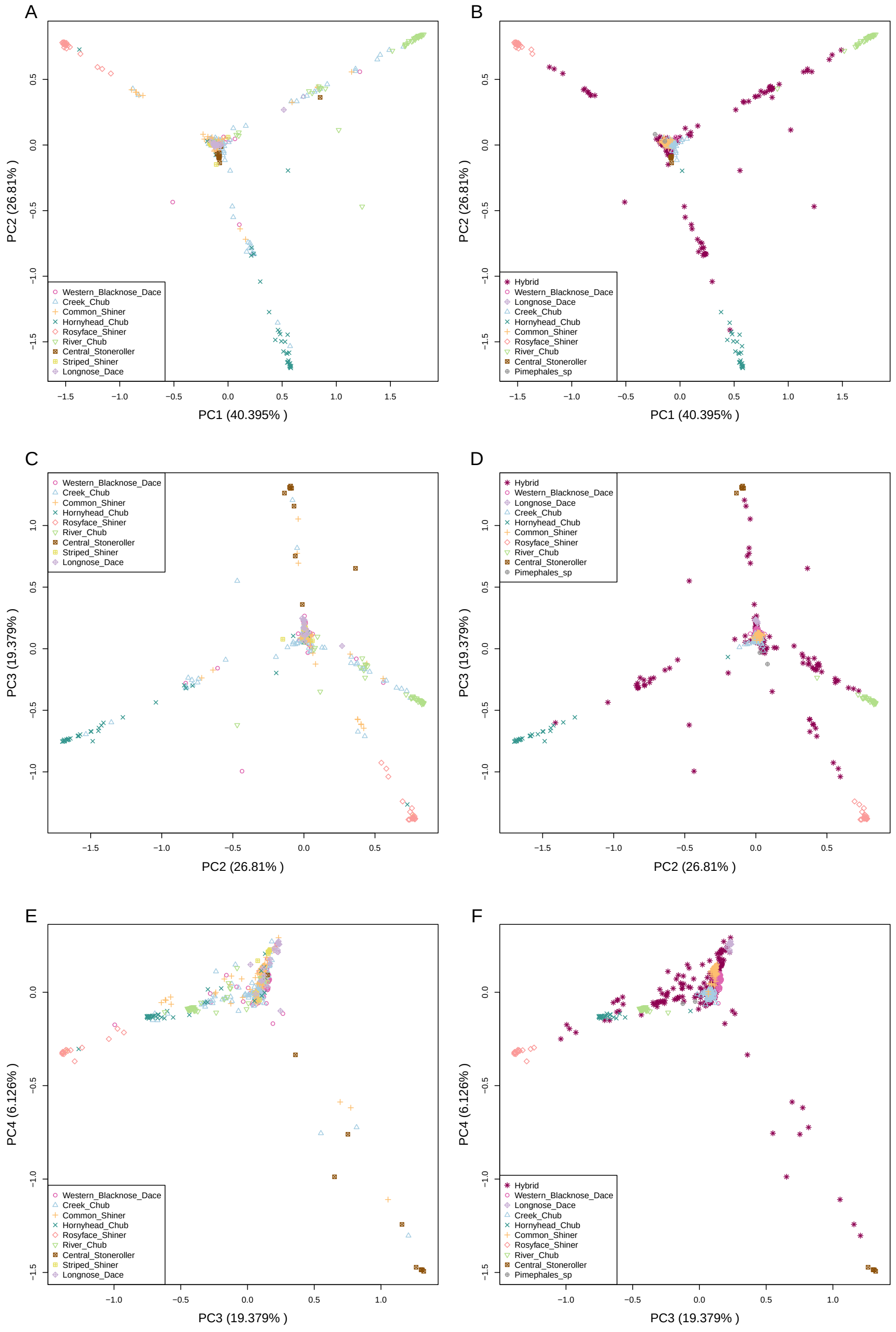


Figure 17S: Principal components 1 through 3 plotted in a step-wise fashion, plotted with both phenotypic and genomic identities. Plots in the left column are coloured by phenotypic identity, while plots in the right column are coloured by genomic identity as determined by `ENTROPY`. A) and B) PC1 and PC2. C) and D) PC2 and PC3. E) and F) PC3 and PC4.

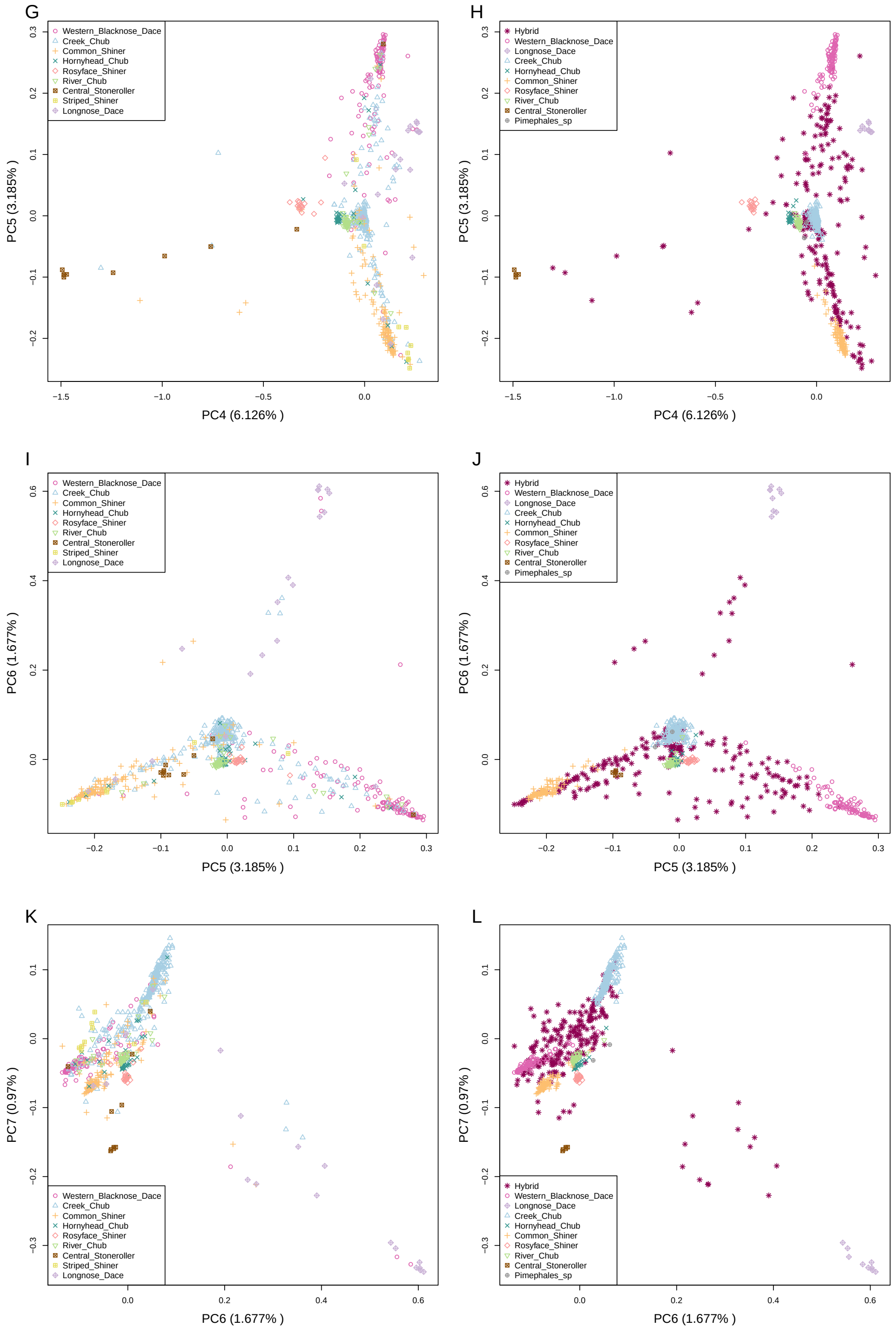


Figure 17S: Principal components 4 through 6 plotted in a step-wise fashion, plotted with both phenotypic and genomic identities. Plots in the left column are coloured by phenotypic identity, while plots in the right column are coloured by genomic identity as determined by `ENTROPY`. G) and H) PC4 and PC5. I) and J) PC5 and PC6. K) and L) PC6 and PC7.
